# Aβ42 oligomers trigger synaptic loss through CAMKK2-AMPK-dependent effectors coordinating mitochondrial fission and mitophagy

**DOI:** 10.1101/637199

**Authors:** Annie Lee, Chandana Kondapalli, Daniel M. Virga, Tommy L. Lewis, So Yeon Koo, Archana Ashok, Georges Mairet-Coello, Sebastien Herzig, Reuben Shaw, Andrew Sproul, Franck Polleux

## Abstract

During the early stages of Alzheimer’s disease (AD) in both mouse models and human patients, soluble forms of Amyloid-β1-42 oligomers (Aβ42o) trigger loss of excitatory synapses (synaptotoxicity) in cortical and hippocampal pyramidal neurons (PNs) prior to the formation of insoluble Aβ plaques. We observed a spatially restricted structural remodeling of mitochondria in the apical tufts of CA1 PNs dendrites in the hAPP^SWE,IND^ transgenic AD mouse model (J20), corresponding to the dendritic domain receiving presynaptic inputs from the entorhinal cortex and where the earliest synaptic loss is detected in vivo. We also observed significant loss of mitochondrial biomass in human neurons derived from a new model of human ES cells where CRISPR-Cas9-mediated genome engineering was used to introduce the ‘Swedish’ mutation bi-allelically (APP^SWE/SWE^). Recent work uncovered that Aβ42o mediates synaptic loss by over-activating the CAMKK2-AMPK kinase dyad, and that AMPK is a central regulator of mitochondria homeostasis in non-neuronal cells. Here, we demonstrate that Aβ42o-dependent over-activation of CAMKK2-AMPK mediates synaptic loss through coordinated MFF-dependent mitochondrial fission and ULK2-dependent mitophagy in dendrites of PNs. We also found that the ability of Aβ42o-dependent mitochondrial remodeling to trigger synaptic loss requires the ability of AMPK to phosphorylate Tau on Serine 262. Our results uncover a unifying stress-response pathway triggered by Aβo and causally linking structural remodeling of dendritic mitochondria to synaptic loss.

## Introduction

Alzheimer’s disease (AD) is the most prevalent form of dementia in humans leading to socially devastating cognitive defects with no effective treatment. AD is characterized by two hallmark lesions including amyloid plaques composed of fibrillar forms of Amyloid-β (Aβ) and neurofibrillary tangles (NFTs) composed of aggregated, hyperphosphorylated Tau. Before the formation of amyloid plaques and NFTs, amyloidogenic processing of amyloid precursor protein (APP) by β- and γ-secretase produces abnormal accumulation of a 42-amino acid-long amyloid beta (Aβ42) peptide. Aβ42 peptides have a strong ability to oligomerize and form dimers, trimers, and higher order oligomers that ultimately fibrillate to form Aβ plaques (Sheng et al., 2012). Oligomeric forms of Aβ42 (Aβ42o) lead to early loss of excitatory synapses (synaptotoxic effects) in cortical and hippocampal pyramidal neurons (PNs) before plaque formation, strongly suggesting that synaptotoxicity is an early event in the disease progression triggered by soluble Aβ42o (Hong et al., 2016; Mairet-Coello et al., 2013; Masliah et al., 2001; Moolman et al., 2004; Shankar et al., 2007). This phenotype has been observed in various AD mouse models including the J20 transgenic mouse model (APP^SWE, IND^) where progressive loss of excitatory synaptic connections occur prior to Aβ plaque formation (Jacobsen et al., 2006; Mairet-Coello et al., 2013; Mucke et al., 2000a). Soluble Aβ42o extracted biochemically from AD patients or produced synthetically also lead to rapid synaptic loss in PNs *in vitro* (Jin et al., 2011; Lacor et al., 2004; Lacor et al., 2007; Mairet-Coello et al., 2013; Shankar et al., 2007; Shankar et al., 2008).

A second phenotype observed in the early stages of AD progression is structural abnormalities of mitochondria. Because of the terminal postmitotic status and extreme degree of compartmentalization of neurons of the central nervous system (CNS), maintaining mitochondrial homeostasis in neurons is highly dependent on the dynamic balance of fission/fusion and efficient autophagic degradation of aged and damaged mitochondria (Ashrafi and Schwarz, 2015; Lewis et al., 2018; Schwarz, 2013). Mitochondrial fission is carried out by a GTPase protein called Drp1 that needs to be recruited to the outer mitochondrial membrane (OMM) by various proteins including MFF, Fis1, MiD49, and MiD51. The fusion machinery includes Mfn1/2 that is involved in fusion of OMM and OPA1 involved in inner mitochondrial membrane fusion (Burte et al., 2015; Chan, 2006; Youle and van der Bliek, 2012). PNs show strikingly compartmentalized morphology of their mitochondria in the somatodendritic versus axonal domains (Lewis et al., 2018). In most cells, mitochondria play critical physiological functions that might differ between dendrites and axons such as ATP production, Ca^2+^ buffering, and/or lipid biosynthesis (Lewis et al., 2013). Recent evidence points to significant structural remodeling of dendritic mitochondria in hippocampal neurons in multiple AD mouse models and AD patients (Burte et al., 2015; Du et al., 2010; Itoh et al., 2013; Wang et al., 2009a; Xie et al., 2013; Zhang et al., 2016), which is one of the earliest affected brain regions in AD in mouse models and AD patients (Arriagada et al., 1992; Busche et al., 2012; David et al., 2005; Eckert et al., 2008; Rhein et al., 2009; Serrano-Pozo et al., 2011). However, the role played by changes in mitochondria structure and/or function in the disease progression *in vivo*, especially in relation to synaptic loss, remains untested. Moreover, the molecular mechanisms causally linking changes in mitochondria structure/function and synaptic loss remain unknown in the context of AD pathophysiology.

Results from our lab as well as others demonstrated that in neurons, Aβ42o over-activates AMP-activated kinase (AMPK) in a Calcium/calmodulin Kinase Kinase protein 2 (CAMKK2)-dependent manner and preventing CAMKK2 or AMPK over-activation either pharmacologically or genetically protects hippocampal neurons from Aβ42o-dependent synaptic loss *in vitro* and in the hAPP^SWE, IND^ transgenic mouse model (J20) *in vivo* (Ma et al., 2014; Mairet-Coello et al., 2013; Son et al., 2012; Thornton et al., 2011). Importantly, AMPK is over-activated in the brain of AD patients where catalytically active AMPK accumulates in pyramidal neurons of the cortex and hippocampus (Fang et al., 2019; Vingtdeux et al., 2011). In non-neuronal cells, AMPK is a key metabolic sensor that is catalytically activated upon increasing levels of AMP/ADP (when ATP levels drop) and regulates various downstream effectors involved in maintaining mitochondrial integrity in part via two recently identified downstream effectors: mitochondrial fission factor (MFF) and Unc-51 like autophagy activating kinases 1/2 (ULK1/2) (Egan et al., 2011; Toyama et al., 2016). MFF is an outer mitochondrial protein with two identified AMPK phosphorylation sites. Upon AMPK activation, MFF enhances the recruitment of the fission protein Drp1, which is important for efficient fragmentation of mitochondria (Toyama et al., 2016), a step thought to be required for efficient mitophagy. The second and coordinated step regulated by AMPK activation are the ULK proteins. ULK1 (also called Atg1) has multiple AMPK phosphorylation sites and functions in an evolutionarily conserved process called autophagy. Autophagy is involved in degrading cytoplasmic components via lysosomes, often under nutrient stress to obtain energy. Importantly, autophagy is the only known mechanism for degrading organelles including mitochondria (mitophagy). Mounting evidence has also implicated altered autophagy in the pathogenesis of AD as abnormal autophagic vacuole (AV) build-up in neurites of AD patients at least at late stages of the disease progression (Nixon, 2013). Therefore, we wanted to test if the CAMKK2-AMPK pathway could be the first pathway identified to coherently regulate the major AD phenotypes and more precisely if Aβ42o-dependent over-activation of AMPK triggers synaptic loss through its ability to mediate mitochondrial structural remodeling.

In the present study, we first reveal that hippocampal CA1 PNs display remarkably compartmentalized structural remodeling of dendritic mitochondria in the J20 AD mouse model *in vivo*: we observe significant loss of mitochondrial biomass only in the apical tuft dendrites of CA1 PNs which, most significantly, corresponds to the dendritic compartment in the hippocampus where loss excitatory synapses can be first observed in this and other AD mouse models (Siskova et al., 2014). We demonstrate that Aβ42o application induces sequential mitochondrial fission and loss of mitochondrial biomass through mitophagy in dendrites of cortical PNs via coordinated phosphorylation and activation of two critical AMPK effectors: MFF and ULK2. Importantly, individual manipulation of either MFF or ULK2 prevents Aβ42o-dependent dendritic spine loss, demonstrating a causal link between Aβ42o-dependent mitochondria structural remodeling and synaptic maintenance. Finally, we previously identified that Aβ42o-dependent AMPK phosphorylates Tau at Serine 262 and that preventing S262 phosphorylation by AMPK protected cortical PNs from Aβ42o-induced synaptic loss. Interestingly, we show that that expression of Tau-S262A is protective towards Aβ42o-induced dendritic mitochondrial remodeling and synaptic loss in cortical PNs. Our results (1) demonstrate that the CAMKK2-AMPK stress-response pathway mediates Aβ42o-dependent synaptic loss through Tau-S262 and MFF-ULK2-dependent remodeling of dendritic mitochondria, (2) provide the first ‘unifying’ stress response pathway triggering many of the cellular defects reported in AD mouse models and in the brain of AD patients during early stages of the disease progression, such as Aβ42o-dependent Tau phosphorylation, defect in Ca^2+^ homeostasis, increased autophagy and defective mitochondrial integrity (reviewed in (Sheng et al., 2012)) and (3) provide the first causal link between Aβ- and Tau-dependent dendritic mitochondrial remodeling and synaptic loss.

## Results

### Spatially restricted structural remodeling of mitochondria and synaptic loss in apical tufts dendrites of CA1 hippocampal PNs in the APP^SWE, IND^ mouse model

The J20 mouse line is a powerful model to study the effect of amyloidosis in AD pathogenesis as it expresses the human Amyloid Precursor Protein (APP) carrying two mutations found in familial forms of AD (APP^SWE, IND^ i.e. KM670.NL and V717F called J20 model thereafter) and show Aβ42o accumulation in the hippocampus (Mucke et al., 2000a). For this study, we chose to focus on the analysis of mitochondrial morphology and dendritic spine density in CA1 hippocampal neurons, which are among the first and the most affected in AD patients and AD mouse models (Perez-Cruz et al., 2011; Pozueta et al., 2013; Serrano-Pozo et al., 2011; Siskova et al., 2014). Using sparse *in utero* hippocampal or cortical electroporation (E15◊P21), we were able to visualize dendritic mitochondria morphology and spine density in optically-isolated dendritic segments of hippocampal CA1 PNs or layer 2/3 pyramidal neurons *in vivo* (Fig. S1). Interestingly, in WT mice, layer 2/3 cortical PNs display elongated and fused dendritic mitochondria, forming a complex network throughout the entire dendritic arbor (Fig. S1B, D, F, H) (Lewis et al., 2018). In sharp contrast, CA1 PNs display a high degree of compartmentalization of mitochondrial morphology in dendrites of WT mice: in basal and apical oblique dendrites, mitochondria are mostly small and punctate whereas the distal apical dendrites contain elongated and fused mitochondria ((Fig. S1A, C, E, G). The transition between these two types of mitochondrial morphology is sharp and corresponds to the boundary between the hippocampal layer stratum radiatum (SR; receiving primarily inputs from CA3 PNs; Fig. 3B) and stratum lacunosum-moleculare (SLM; receiving exclusively presynaptic inputs from the medial entorhinal cortex MEC; Fig. 3B).

We applied the same experimental approach in both WT and J20 littermate mice and examined mitochondrial morphology and spine density in 3-month-old mice, a time-point where abundant Aβ42o can be detected but preceding appearance of amyloid plaques (Mucke et al., 2000a). This analysis revealed that, consistent with what we observed in hippocampus of WT mice at P21 (Fig. S1), 3 month-old WT and J20 mice both display mostly small, punctate mitochondria in basal and apical oblique dendrites (Fig. 1A). This is accompanied by a trend but not significant decrease in spine density in basal and apical oblique dendrites (Fig. 1A, C). However, the distal apical dendrites of the J20 mice showed a dramatic and significant reduction of both mitochondria length and density/biomass as well as a significant decreased in spine density compared to WT littermates (Fig. 1A, C, E). These results reveal (1) a striking and unique degree of compartmentalization of mitochondrial morphology in dendrites of CA1 PNs not observed in cortical layer 2/3 PNs and (2) a spatially restricted loss of mitochondrial biomass and spine density specifically in the apical tuft dendrites of CA1 PNs in the J20 mouse model. This co-occurrence of mitochondrial remodeling and decreased spine density raised the possibility, tested below, of a potential causal relationship between Aβ42o-dependent mitochondrial remodeling and synaptic maintenance.

**Figure 1.**
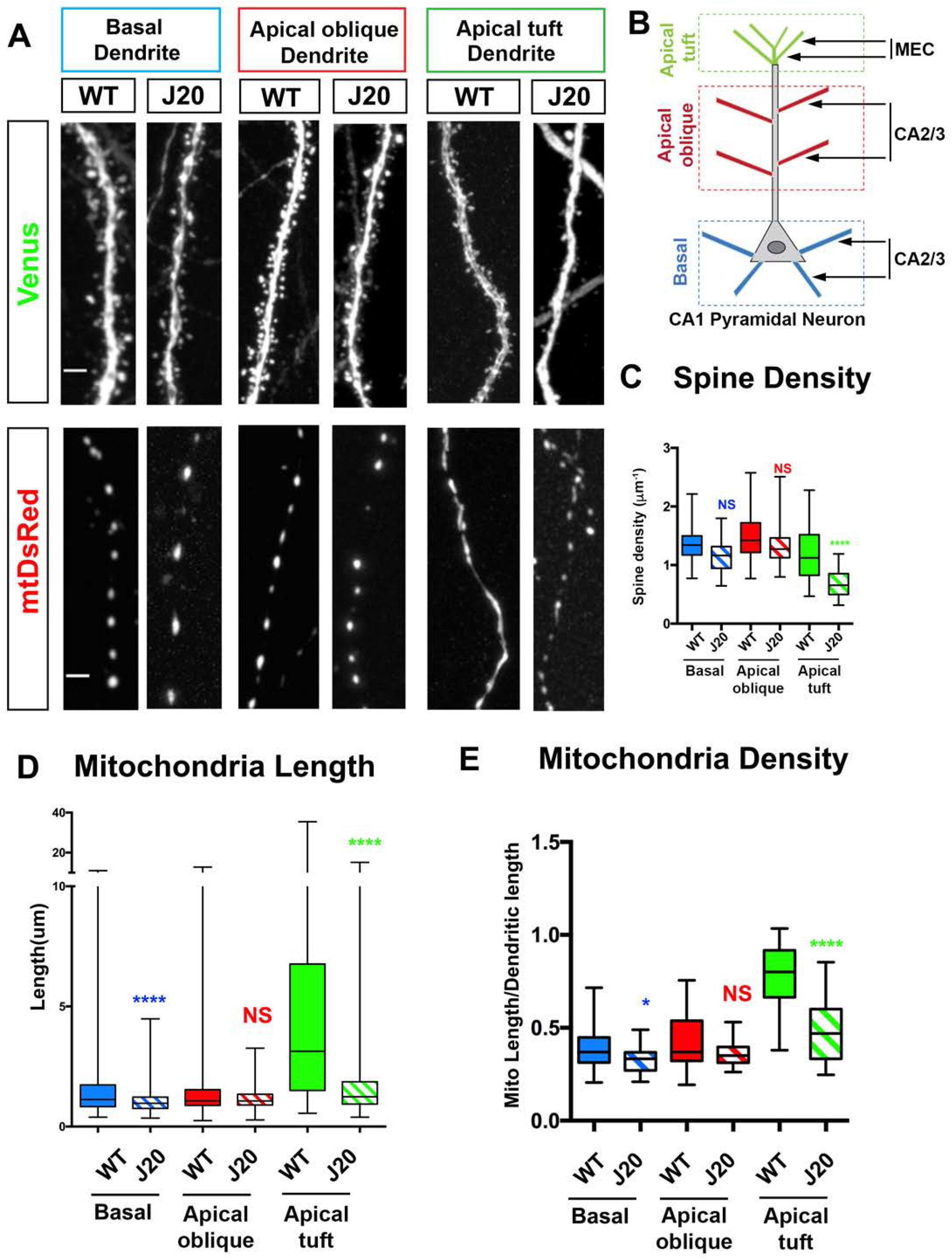
Aβ42o induces dendritic mitochondrial fragmentation and dendritic spine loss within the same time frame. (A-C’) Representative images of primary cortical neurons at 21 days *in vitro* (DIVs). Neurons express Venus (A,B,C) and mito-DsRed (A’, B’, C’) to assess spine density and mitochondrial morphology, respectively. At 20 DIVs, neurons were treated with either 24 hours of control vehicle, 14 or 24 hours of Aβ42o (300-450nM-see Material and Methods for details). High magnification of secondary dendrites is shown below whole neurons shown at low magnification. (D) Quantification of dendritic mitochondrial length. (E) Quantification of dendritic mitochondria density. Dendritic mitochondria density was calculated by summing up the length of all the mitochondria then dividing it by the length of the dendrite examined. (F) Quantification of dendritic spine density calculated by dividing the number of spines by the length of the dendrite examined. All the analysis was done blind to the experimental conditions, and was done by manual counting using FIJI. All of the statistical analyses were performed with Prism 6 (GraphPad Software). Data is represented by box plots displaying minimum to maximum values, with the box denoting 25^th^, 50^th^ (median) and 75^th^ percentile from 3 independent experiments. n_control_ = 53 dendrites, 278 mitochondria; n_14hr Aβ42_ = 47 dendrites; 380 mitochondria; n_24hr Aβ42=_ 51 dendrites; 400 mitochondria. Statistical analysis was performed using Kruskal-Wallis test followed by Dunn’s post-hoc test in (D-F). The test was considered significant when p<0.05 with the following criteria: * p<0.05; ** p<0.01; ***p<0.001; ****p<0.0001; ns, not significant. Scale bar for high magnification dendritic segments= 2μm.

### Aβ42o induces synaptic loss and dendritic mitochondrial remodeling in the same time frame

In order to examine the temporal relationship between mitochondrial remodeling and decreased spine density, we wanted to adopt an in vitro model that would allow us to control the timing of Aβ42o application. We turned to long-term cortical PNs cultures (Mairet-Coello et al., 2013) which are synaptically connected and display uniformly fused mitochondria throughout their dendritic arbor *in vitro* (Fig. 2) as observed *in vivo* (Fig. 1 and S1). We observed significant, time-dependent changes in dendritic mitochondrial structure following acute Aβ42o application including both a reduction of their length (potentially through increased fission) and reduced mitochondrial density/biomass (Fig. 2A’-C’; Fig. 2D-E). In the same neurons, we observed a progressive reduction in dendritic spine density in Aβ42o treated neurons between 14 and 24hr following Aβ42o application (Fig. 2A-C, Fig. 2F). No significant changes in mitochondria morphology were observed in the soma or the axon of the same neurons (data not shown) or, as previously reported, when control inverted peptide (INV42) were applied at similar concentrations than Aβ42o (300-450nM; Fig. S2E) (Lacor et al., 2004; Lacor et al., 2007; Mairet-Coello et al., 2013; Shankar et al., 2007). We previously published that at this dose and duration, Aβ42o application does not affect neuronal viability, strongly suggesting that the effects we observe are not a secondary consequence of compromising neuronal survival (Mairet-Coello et al., 2013). Taken together, our results highlight that Aβ42o application triggers a significant degree of time-dependent, structural remodeling of dendritic mitochondria including reduction of mitochondrial biomass concomitant with a reduction in spine density, which is a reliable index of excitatory synapse number in dendrites of PNs (Harnett et al., 2012).

**Figure 2.**
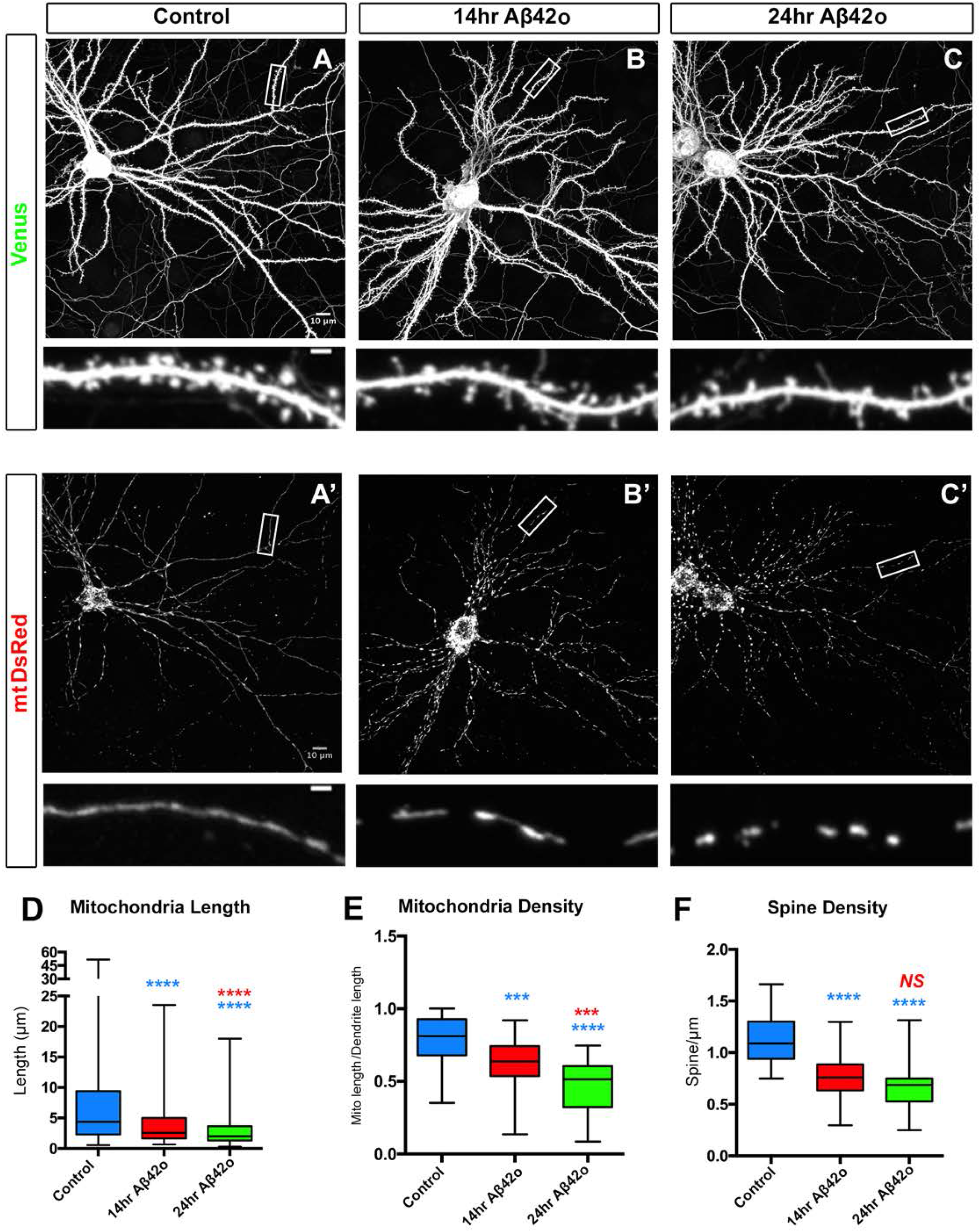
Aβ42o treatment induces local mitophagy in dendrites. **(A)** Dendritic segments (secondary) from primary cortical PNs at 21-25 DIVs electroporated with plasmids encoding mito-mTagBFP2, Lamp1-mEmerald, and RFP-LC3 to visualize mitochondria, lysosomes, and autophagosomes, respectively, using *ex utero* electroporation on embryos at E15.5. At 21DIVs, neurons were treated with control vehicle or Aβ42o and imaged live with time-lapse microscopy every 15 minutes for 14 hours. The captured image shows dendrites at 0 and 14 hours for both conditions. Yellow box labels the zoomed region shown in **(A’)** illustrating that in control, there is no significant change of LAMP1+ (lysosomes) and LC3+ (autophagosomes) or LC3/LAMP1 double-positive (autolysosomes) vesicles dynamics or accumulation and no significant change in mitochondrial morphology (see also Movie S1 and S2). In contrast, in dendrites of Aβ42o treated cortical PNs, sites of LAMP1-LC3 double-positive autolysosome accumulation often results in loss of mitochondria as indicated by yellow arrow in A’. We note that some sites do not display loss of mitochondrial signal as within the 14h of imaging following Aβ42o application (red arrow in Fig. 2A). **(A’’)** Representative kymograph of mitochondria, lysosomes, and autophagosomes in dendrites from Fig. 2A. **(B)** Percentage of dendritic segments showing accumulation of both LC3+ autophagosomes, LAMP1+ lysosomes or LC3/ LAMP1-double positive autolysosomes. Data is represented by box plots displaying minimum to maximum values, with the box denoting 25^th^, 50^th^ (median) and 75^th^ percentile from 3 independent experiments. n_control_ = 15 neurons; n_Aβ42_ = 24 neurons. **(C)** Random dendritic segments analyzed in B were selected to measure levels of mitophagy. To obtain a spatially specific measurement of mitophagy, intensity line scan measurements were taken across different dendritic regions from the same dendrite. For each dendrite, line scans were drawn in regions without and with LC3 and LAMP1 puncta at 14 hours and the intensity of mitochondrial signal was measured at time point zero and 14 hours. Fold change of mitochondrial intensity at a very spatially focused area was calculated. In control, mitochondrial index across the same dendrite is relatively unchanged in regions with and without LC3 and LAMP1 accumulation, suggesting that mitochondria are not targeted for LC3/LAMP1 degradation. In Aβ42 treated condition, mitochondrial index along the same dendrite can show significant difference. Only regions of LC3 and LAMP1 accumulation show mitochondrial loss. Data is represented as bar graph from n _control conditions_ = 33 dendrites; n _Aβ42 conditions_ = 44 dendrites. One-way ANOVA with Kruskal-Wallis followed by Dunn’s post-hoc test were applied. The test was considered significant when p<0.05 with the following criteria: * p<0.05; ** p<0.01; ***p<0.001; ****p<0.0001; ns, not significant. Scale bar= 2μm.

These results show that *in vivo* exposure of CA1 hippocampal PNs to Aβ42o resulting from over-expression of APP^SWE,IND^ in the J20 AD mouse model or direct application of synthetic Aβ42o at 300-450 nM for 24h in mature cortical PNs *in vitro* leads to striking loss of dendritic mitochondrial biomass and synaptotoxicity. However, two potential criticisms of these approaches are (1) that they both use mouse models which could differ from the way human neurons could respond and (2) that both approaches rely on exposure of cortical or hippocampal PNs to high levels of Aβ42o. In order to address both points, we generated a novel human embryonic stem cell (hESC) APP^SWE/SWE^ knockin model. CRISPR-Cas9-mediated genome engineering was used to introduce to introduce the APP^SWE^ mutation into both alleles of the H9 hESC cell line (WA09, WiCell). Previous knockin of the Swedish mutation into a human PSC line demonstrated increased endogenous APP processing and generation of Aβ42o in an allelic dose-dependent manner (Paquet et al., 2016).

APP^SWE^ knockin hESCs and their isogenic H9 parent control line were transdifferentiated into glutamatergic cortical neurons (iNs) using doxycycline-inducible Neurogenin2 (Ngn2) expression (Zhang et al., 2013). Following efficient Ngn2-P2A-eGFP-induced neuronal differentiation (see Methods), we transiently transfected these neurons in a sparse manner with a mitochondrial fluorescent reporter (mitoDsRed) and quantified dendritic mitochondrial morphology in control H9 and isogenic APP^SWE/SWE^-derived neurons (Fig. S3A). Our results demonstrate a significant reduction in dendritic mitochondrial density (Fig. S3B) and dendritic mitochondrial length (Fig. S2C) in APP^SWE/SWE^ human neurons compared to isogenic control neurons. As previously observed in cortical and hippocampal neurons in vitro and in vivo (Mairet-Coello et al., 2013), we also found increased AMPK activation in APP^SWE/SWE^ neurons compared to isogenic control neurons, as measured biochemically by the ratio of phosphorylated AMPK-T172 to total AMPK (Fig. S3D-E). These results demonstrate that endogenous levels of Amyloid-*β* generated by processing of APP in human mutant neurons can trigger dendritic mitochondrial remodeling and loss of biomass as well as AMPK over-activation.

### Aβ42o application induces local mitophagy in dendrites

We next tested whether changes in mitochondrial biomass observed upon Aβ42o treatment is triggered in dendrites of PNs by local mitophagy. To that end, we examined the dynamics of autophagosomes, lysosomes, autolysosomes as well as changes in mitochondrial biomass in cortical PNs upon Aβ42o treatment using sparse *ex utero* electroporation of plasmids encoding mito-mTagBFP2, LAMP1-mEmerald, and RFP-LC3 to visualize mitochondria, lysosomes, and autophagosomes, respectively. Cortical PNs were cultured for at least 21DIV in high-density cultures, where they display mature synapses (Mairet-Coello et al., 2013), and imaged live using time-lapse confocal microscopy every 15 minutes for 14 hours following treatment with either control (vehicle) or Aβ42o (Fig. 3). In control treated neurons, we observed little to no change in mitochondrial biomass or in the dynamics of LC3+ (autophagosomes), LAMP1+ (lysosomes) or LC3+/LAMP1+ (autolysosomes) vesicles (Fig. 3A-A’’). On the contrary, upon Aβ42o treatment, we observed a significant accumulation of both LC3+ and LAMP1+ puncta in dendrites over time as visualized in Fig. 3A (see kymograph in Fig. 3A”). In dendrites exposed to Aβ42o, many of the LC3+ puncta became co-localized with LAMP1+, suggesting these autolysosomes often occur locally. LC3+/LAMP1+ autolysosomes do form occasionally in control dendrites (Fig. 3A’’-B). As shown in Fig. 3B, treatment with Aβ42o leads to an almost 5-fold increase in the percentage of dendritic segments with increased LC3+/LAMP1+ autolysosomes accumulation compared to control treatment. Both LC3 and LAMP1 normalized fluorescence intensity showed time-dependent increases upon Aβ42o application, highlighting local biogenesis of autophagic vacuoles and their coalescence with lysosomes in dendritic shafts upon Aβ42o treatment, respectively (Fig. 2C-E). We also observed a significant reduction in mitochondria fluorescence (as measured via mt-mTAGBFP2, (Fig. 2E)) in a spatially restricted area close to LC3+/LAMP1+ puncta as indicated by the yellow arrow in the representative image (Fig. 3A) and the kymograph (Fig. 3A”). Not all LC3+/LAMP1+ puncta correlated with loss of a mitochondria within 14h of time-lapse imaging as indicated by the red arrow in the representative image (Fig. 3A-A”). To quantify spatial dynamics of mitophagy, we defined a ‘mitophagy index’ along segments of dendrites by measuring the intensity of mito-mTagBFP2 fluorescence using line scan intensity measurements in regions of interest (ROI) with or without LC3+/LAMP1+ puncta. This index reflects the probability of loss of mitochondrial biomass in or out of areas where autophagosomes and lysosomes coalesce to generate autolysosomes. In control conditions, the formation of LC3+/LAMP1+ puncta is not only a rare event (Fig. 3A-B), but the presence of LC3+/LAMP1+ puncta do not result in loss of mitochondria biomass therefore suggesting that basal levels of mitophagy are exceedingly low in dendrites of cortical PNs in these conditions (Fig. 3B-C). In contrast, upon Aβ42o treatment, we frequently observed spatially restricted mitophagy events, only found in ROIs of the dendrites that display LC3+/LAMP1+ autolysosomes demonstrating that significant levels of mitophagy occur locally in dendrites of cortical PNs upon Aβ42o treatment (Fig. 3B-C).

**Figure 3.**
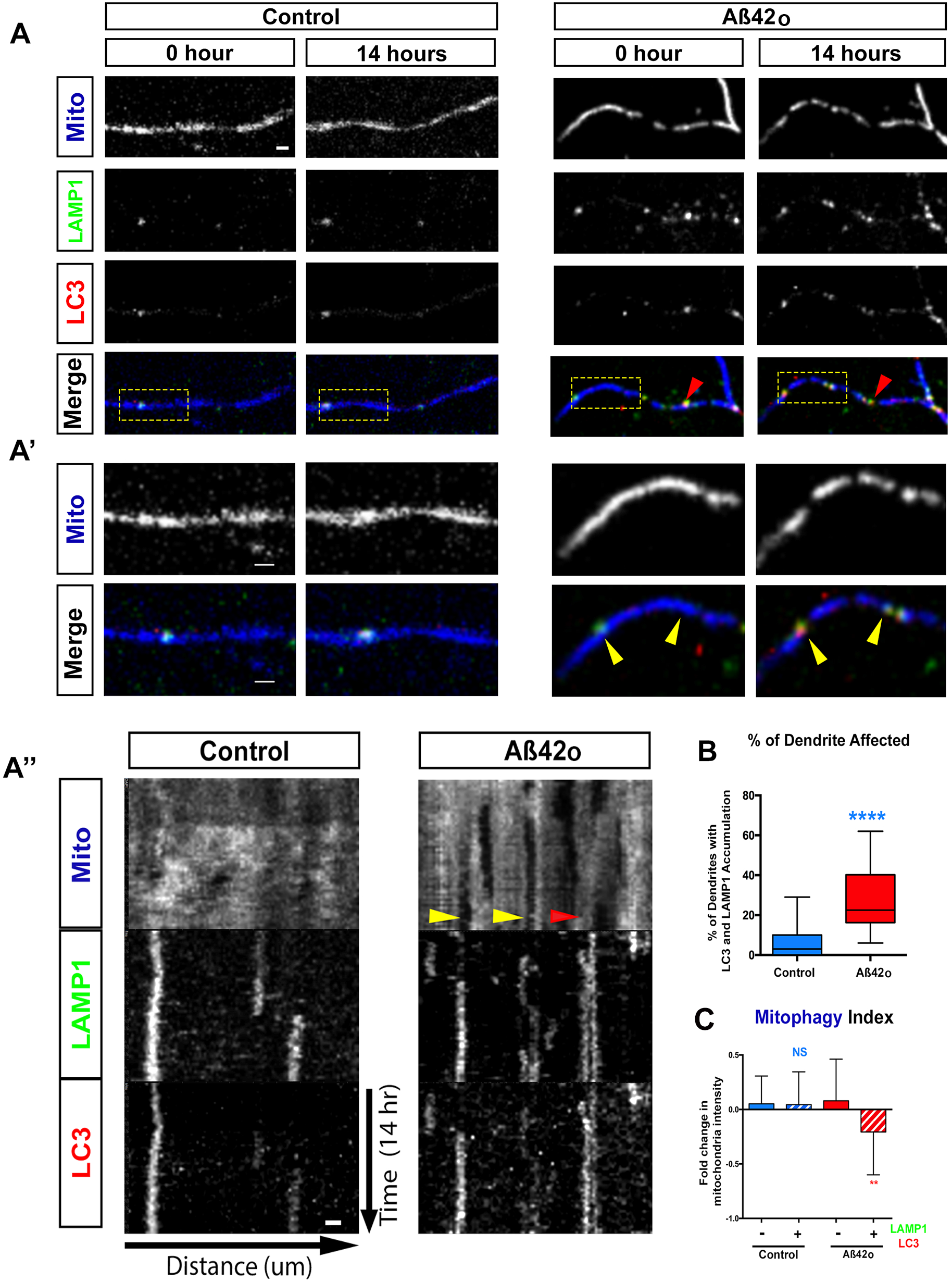
In AD mouse model, CA1 pyramidal neurons display spatially restricted mitochondrial fragmentation and loss of biomass in distal apical tufts. (**A**) Representative images of CA1 PN dendrites from different regions of WT control and J20 mice. CA1 PNs were electroporated with pCAG-Venus and pCAG-mito-DsRed by *in utero* electroporation in E15.5 WT and J20 mouse embryos. Both mouse groups were fixed and imaged at 3 months postnatal (P90). Dendrites from basal, oblique, and distal apical are magnified to show spine density and mitochondrial morphology. The three types of dendrites were characterized as displayed in (**B**) for both WT and J20 mice. (**C-E**) Quantification of spine density (C) and individual mitochondrial length (D) and mitochondrial density (fraction of dendritic segment length occupied by mitochondria) (E). Data is represented by box plots displaying minimum to maximum values, with the box denoting 25^th^, 50^th^ (median) and 75^th^ percentile from at least 3 independent *in utero* electroporated mice. N_basal WT_ = 47 dendrites, 595 mitochondria; N_basal APP_ = 32 dendrites, 495 mitochondria; N_oblique WT_ = 46 dendrites, 630 mitochondria; N_oblique APP_ = 33 dendrites, 507 mitochondria; N_distal WT_ = 39 dendrites, 284 mitochondria; N_distal APP_ = 32 dendrites; 426 mitochondria. All the analysis was done blind to the experimental conditions, and was manually counted using FIJI. Statistical analysis was performed using Kruskal-Wallis test followed by Dunn’s post-hoc test in (C-E). The test was considered significant when p<0.05 with the following criteria: * p<0.05; ** p<0.01; ***p<0.001; ****p<0.0001; ns, p>0.05. Scale bar= 2μm.

### Aβ42o-dependent mitochondrial fragmentation and synaptotoxicity are mediated by CAMKK2-AMPK over-activation

We have previously shown that Aβ42o-dependent over-activation of the CAMKK2-AMPK kinase dyad mediates excitatory synaptic loss both in primary cortical PNs culture and in hippocampal PNs in the J20 AD mouse model, (Mairet-Coello et al., 2013). Since AMPK is a central regulator of mitochondrial homeostasis (Garcia and Shaw, 2017; Herzig and Shaw, 2018), we wanted to test if the CAMKK2-AMPK pathway was required for Aβ42o-induced dendritic mitochondrial remodeling in primary cortical PNs cultures *in vitro* using a conditional double floxed mouse line (AMPKα1^F/F^/α2^F/F^). The CA1 hippocampal region of AMPKα1^F/F^/α2^F/F^ embryos was electroporated at E15.5 with (1) a control plasmid lacking Cre recombinase or a plasmid expressing Cre recombinase along (2) with Venus fluorescent protein to visualize spine morphology and (3) mt-dsRed to examine mitochondrial morphology. Quantitative analysis indicated that in basal conditions AMPKα1/α2 null neurons have spine and mitochondria densities indistinguishable from wildtype neurons. Strikingly, AMPKα1/α2 null neurons are completely protected from Aβ42o-induced mitochondrial density reduction (Fig. 4A) as well as synaptotoxicity as previously reported (Mairet-Coello et al., 2013). This demonstrates that over-activation of AMPK is required for both Aβ42o-induced mitochondria structural remodeling and spine loss in dendrites of PNs. AMPK has two upstream regulatory kinases namely, LKB1 and CAMKK2 which phosphorylates AMPK*α* subunit on T172 and significantly increases its catalytic activity (Garcia and Shaw, 2017). However, in neurons, CAMKK2 has been identified as the primary upstream regulator of AMPK (Hardie et al., 2012; Mairet-Coello et al., 2013; Son et al., 2012). Therefore, we assessed if pharmacologically blocking CAMKK2 acutely interfered with Aβ42o effects on mitochondrial structural remodeling and dendritic spine loss. Pretreatment of primary neuronal cultures with STO-609, a specific CAMKK2 inhibitor (Tokumitsu et al., 2002), completely blocked Aβ42o-induced mitochondrial structural remodeling (Fig. S4A-C) as well as synaptotoxicity (Fig S4A; S4D) as previously published (Mairet-Coello et al., 2013).

**Figure 4.**
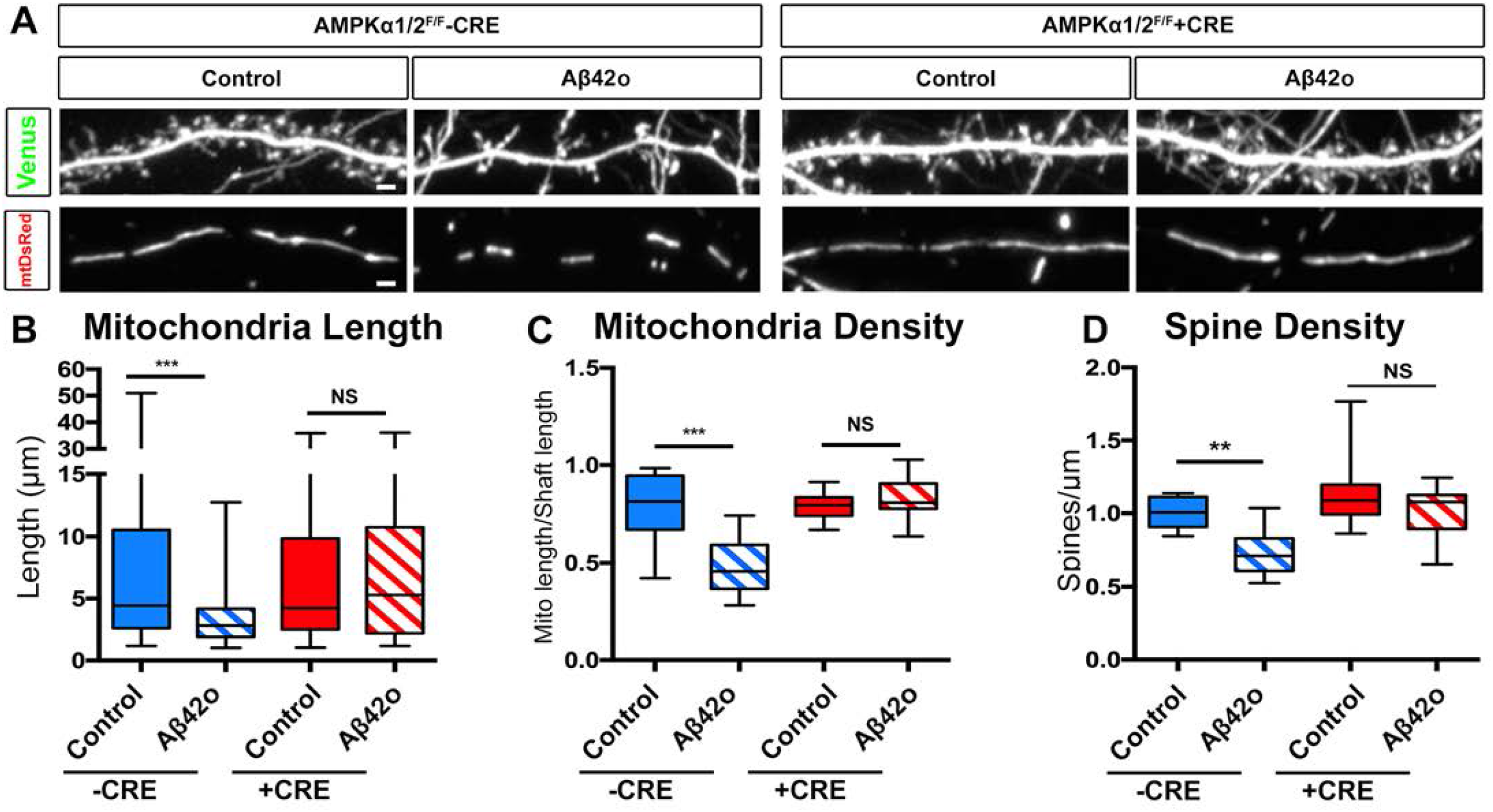
Oligomeric Aβ42 induced synaptotoxicity and dendritic mitochondrial fragmentation is AMPK dependent. (**A**) Dendritic segments (secondary) of primary cortical neuron at 21 DIV. Embryos from AMPKα1/2 double conditional knockout (AMPKα1^F/F^/2^F/F^) were *ex utero* electroporated at E15.5 with plasmids expressing Venus, mito-DsRed, and either scrambled Cre (control) or Cre recombinase. Neurons were treated at 20 DIV with either control or Aβ42o for 24 hr. (**B-D**) Quantification of mitochondria length (**B**), and mitochondrial density (**C**), and spine density (**D**) in individual dendritic segments. All the analysis was done blind to the experimental conditions, and was done by manual counting using FIJI. All of the statistical analyses were performed with Prism 6 (GraphPad Software). Data is represented by box plots displaying minimum to maximum values, with the box denoting 25^th^, 50^th^ (median) and 75^th^ percentile from 3 independent experiments. n_-Cre control_ = 27 dendrites, 128 mitochondria; n_-Cre Aβ42o_ = 29 dendrites; 204 mitochondria; n_+Cre control_ = 29 dendrites; 143 mitochondria; n_+Cre Aβ42o_ =24 dendrites, 112 mitochondria. Statistical analysis was performed using Kruskal-Wallis test followed by Dunn’s post-hoc test in (B-D). The test was considered significant when p<0.05 with the following criteria: * p<0.05; ** p<0.01; ***p<0.001; ****p<0.0001; ns, not significant. Scale bar= 2μm.

### Aβ42o-induced MFF over-activation causally links mitochondrial fragmentation and synaptic loss

In non-neuronal cells, AMPK has been identified as a major metabolic sensor regulating ATP homeostasis upon induction of metabolic stress through its ability to phosphorylate many downstream effectors involved in various aspects of metabolic homeostasis (Hardie et al., 2012; Herzig and Shaw, 2018). As part of this metabolic stress response, AMPK regulates mitochondrial fission through direct phosphorylation of mitochondrial fission factor (MFF) which enhances its ability to recruit Drp1 to the outer mitochondria and promotes mitochondrial fission (Toyama et al., 2016). To that end, we tested if MFF is the effector mediating AMPK-dependent Aβ42o-induced mitochondrial fission and if blocking MFF-dependent fission can prevent synaptic loss. Upon exposure of Aβ42o, cortical PNs *in vitro* that express MFF shRNA (knockdown efficacy shown in Lewis et al., 2018) are protected from loss of mitochondrial biomass in dendrites (Fig. 5A-C) and from synaptic loss (Fig. 5A and 5D). Interestingly, knockdown of MFF in basal condition does not affect either the dendritic mitochondrial length and density or dendritic spine density, strongly suggesting that the signaling pathway we have identified is a *bona fide* stress response pathway in dendrites upon Aβ42o exposure, as we have previously hypothesized (Mairet-Coello and Polleux, 2014). Altogether, these results suggest a causal relationship between Aβ42o-induced MFF-dependent dendritic mitochondria fission and dendritic spine loss.

**Figure 5.**
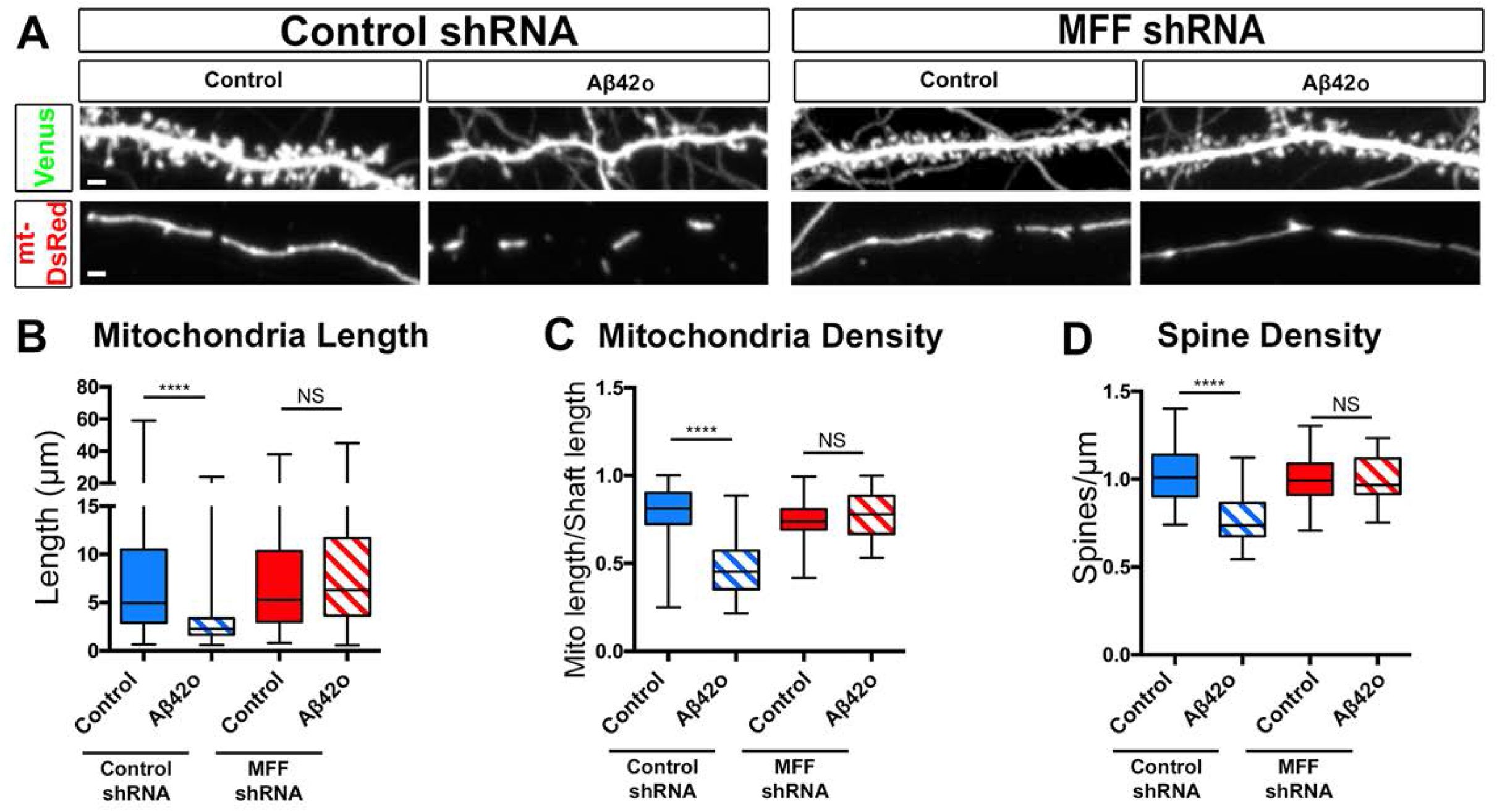
MFF is required for Aβ42o induced dendritic mitochondrial fragmentation. (**A**) Representative images of secondary dendritic segments of primary cortical neurons at 21 DIV. Embryos at E15.5 were *ex utero* electroporated with plasmids encoding Venus, mito-DsRed, and either with control shRNA or an shRNA specific for mouse MFF (MFF shRNA). Neurons were treated at 20 DIV with either control or Aβ42o for 24 hours. Knockdown of MFF blocks both the synapse and dendritic mitochondrial fragmentation. (**B**) Quantification of dendritic mitochondrial length, and (**C**) dendritic mitochondrial density, and (**D)** spine density. All the analysis was done blind to the experimental conditions, and was done by manual counting using FIJI. All of the statistical analyses were performed with Prism 6 (GraphPad Software). Data is represented by box plots displaying minimum to maximum values, with the box denoting 25^th^, 50^th^ (median) and 75^th^ percentile from 3 independent experiments. n_PLKO control_ = 36 dendrites, 188 mitochondria; n_PLKO Aβ42o_ = 41 dendrites; 300 mitochondria; n_MFFshRNA control_ = 36 dendrites; 167 mitochondria; n_MFFshRNA Aβ42o_ = 30 dendrites, 125 mitochondria. Statistical analysis was performed using Kruskal-Wallis test followed by Dunn’s post-hoc test in (B-D). The test was considered significant when p<0.05 with the following criteria: * p<0.05; ** p<0.01; ***p<0.001; ****p<0.0001; ns, not significant. Scale bar for magnified dendritic segments= 2μm.

### Aβ42o-induced MFF phosphorylation by AMPK is required for dendritic mitochondrial fragmentation and synaptic loss

To determine if MFF is phosphorylated by Aβ42o-dependent AMPK over-activation in neurons, we measured AMPK-mediated phosphorylation on MFF at Serine 146 (S146; one of the AMPK phosphorylation site of Isoform 3 enriched in the brain; (Ducommun et al., 2015)) and total MFF in primary cortical neurons (Fig. S5). As a positive control to show that the MFF S146 site is regulated by AMPK activation, we over-expressed the human isoform of MFF in Neuro2A (N2A) neuroblastoma cells and treated with or without Metformin, a potent activator of AMPK (Toyama et al., 2016) (Fig. S5). Firstly, Aβ42o application for 24h on long-term cortical PNs cultures showed a significant increase in AMPK phosphorylation at T172 site as previously reported (Mairet-Coello et al., 2013; Thornton et al., 2011) (Fig. S5). As a consequence of AMPK activation, we observe an increase in phosphorylation of MFF at S146 in cortical neurons upon Aβ42o application which is comparable to Metformin treatment of N2A cells (Fig. S5). This experiment demonstrates that Aβ42o leads to AMPK dependent phosphorylation and activation of MFF in cortical PNs.

In order to test if AMPK-dependent phosphorylation of MFF is required for Aβ42o-dependent mitochondrial remodeling, we took a gene replacement approach where we identified low expression levels of wild-type human MFF (*hMFF WT* not targeted by shRNA directed against mouse MFF) and hMFF carrying two point mutations (S155A and S172A referred to as *hMFF AA* previously shown to abolish AMPK-mediated MFF function in mitochondrial fission; (Toyama et al., 2016)) that, when combined with mouse MFF shRNA, does not lead to mitochondrial fragmentation in control conditions i.e. does not lead to MF over-expression which was previously shown to be sufficient to induce dendritic mitochondrial fragmentation (Toyama et al., 2016). Primary cortical PNs expressing these titrated levels of hMFF WT showed both mitochondrial and spine loss following 24hr of Aβ42o treatment (Fig. 6C-F) to levels comparable to control neurons exposed to Aβ42o, whereas neurons expressing titrated levels of hMFF AA were protected from Aβ42o-induced mitochondrial structural remodeling and spine loss (Fig. 6C-F). These results demonstrate a causal role of Aβ42o-triggered mitochondrial remodeling induced by AMPK-mediated phosphorylation of MFF and synaptic loss.

**Figure 6.**
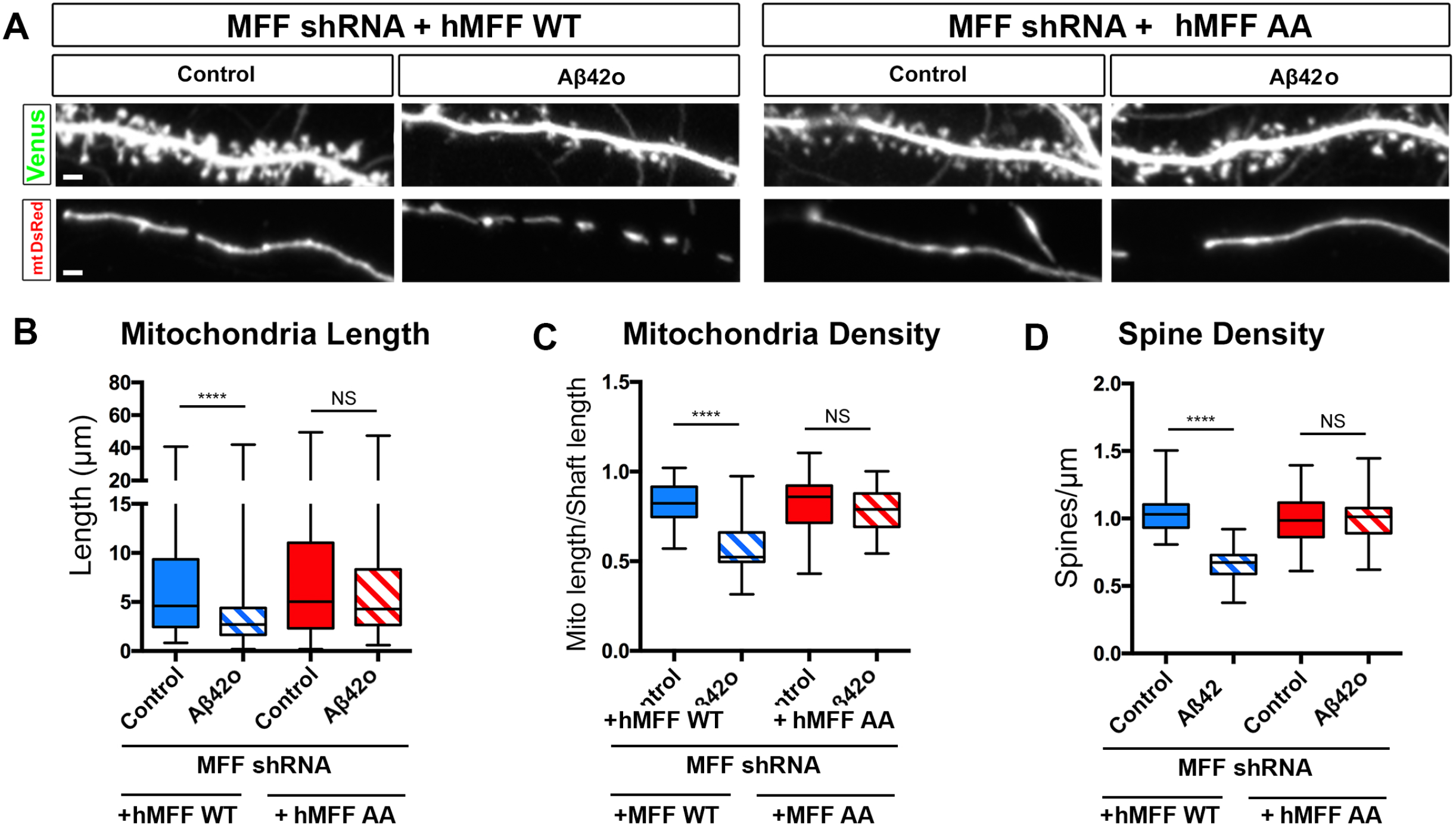
Aβ42o induces AMPK-dependent MFF phosphorylation at two serine sites required for Aβ42o-dependent dendritic mitochondrial fragmentation and spine loss. (**A**) Representative images of secondary dendritic segments of primary cortical neurons at 21 DIVs. Embryos at E15.5 were *ex utero* electroporated with pCAG-Venus and pCAG-mito-DsRed to visualize spines and mitochondria, respectively. Neurons were also electroporated with MFFshRNA along with either low levels of pCAG-MFF-WT or pCAG-MFF-AA mutant constructs and treated with either control or Aβ42o for 24 hours. Gene replacement (MFFshRNA + MFF AA) with a form of MFF that cannot be phosphorylated by AMPK blocks dendritic mitochondrial fragmentation and spine loss. (**B**) Quantification of dendritic mitochondrial length, and (**C**) dendritic mitochondrial density, and (**D**) spine density. All the analysis were done blind to the experimental conditions, and performed by manual counting using FIJI. Data is represented by box plots displaying minimum to maximum values, with the box denoting 25^th^, 50^th^ (median) and 75^th^ percentiles from 3 independent experiments. n_MFFshRNA + MFFWT Control_ = 31 dendrites, 166 mitochondria; n_MFFshRNA + MFFWT Aβ42o_ = 29 dendrites; 228 mitochondria; n_MFFshRNA + MFFAA control_ = 35 dendrites; 164 mitochondria; n_MFFshRNA + MFFAA Aβ42o_ =28 dendrites, 137 mitochondria. Statistical analysis was performed using Kruskal-Wallis test followed by Dunn’s post-hoc test in (D-F). The test was considered significant when p<0.05 with the following criteria: * p<0.05; ** p<0.01; ***p<0.001; ****p<0.0001; ns, not significant. Scale bar in A: 2μm.

### Activation of ULK2 by AMPK leads to loss of mitochondrial biomass following MFF-dependent mitochondrial fission

In non-neuronal cells under conditions of metabolic stress, AMPK activation not only triggers increased mitochondrial fission through phosphorylation and activation but also trigger mitophagy by directly phosphorylating the autophagy initiating kinase, ULK1 (Atg1). Since we observed (1) increased mitophagy in dendrites exposed to Aβ42o and (2) a significant decrease in mitochondrial biomass following increased AMPK-MFF-dependent fission between 14-24h of Aβ42o treatment (Fig. 1 and 5), we hypothesized that over-activation of AMPK by Aβ42o also activates the observed increased levels of mitophagy by activating ULK1 and/or ULK2, two isoforms of ULK implicated in mitophagy and highly expressed in cortical neurons (Egan et al., 2011; Kim et al., 2011).

We tested first if either one or both of the two isoforms of the ULK1/2 mediate Aβ42o-induced synaptotoxicity by performing shRNA-mediated knockdown of either ULK1 or ULK2 (Fig. S6A). Interestingly, we found that knockdown ULK2, but not ULK1, protected cortical PNs from Aβ42o-induced spine loss at 24h (Fig. S6B-C). We then tested if loss of ULK2 could protect Aβ42o-treated neurons from loss of mitochondrial biomass. We sparsely expressed Cre-recombinase in primary cortical PNs isolated from a conditional ULK2 mouse line (ULK2^F/F^) (Cheong et al., 2011) along with Venus as a cell filler and mt-dsRed to visualize mitochondria. Loss of ULK2 completely blocks Aβ42o-induced loss of mitochondrial biomass, however, interestingly, dendritic mitochondria remain fragmented compared to the controls (Fig. 7A-C), compatible with the AMPK-MFF dependent induction of mitochondrial fission being intact in these ULK2-deficient neurons. This further supports the current idea that mitochondrial fission and mitophagy are coupled events such that mitochondrial fragmentation must precede mitophagy (Chan, 2006; Schwarz, 2013). Interestingly, preservation of mitochondrial density/biomass upon ULK2 conditional deletion was sufficient to rescue Aβ42o-induced synaptotoxicity (Fig.7A and D). Therefore, our data suggest that these two parallel effectors (MFF and ULK2) coincidentally activated by AMPK act in a concerted manner to couple fission and mitophagy in neuronal dendrites and most importantly that loss of mitochondrial biomass triggered by Aβ42o is causally linked to synaptic loss.

**Figure 7.**
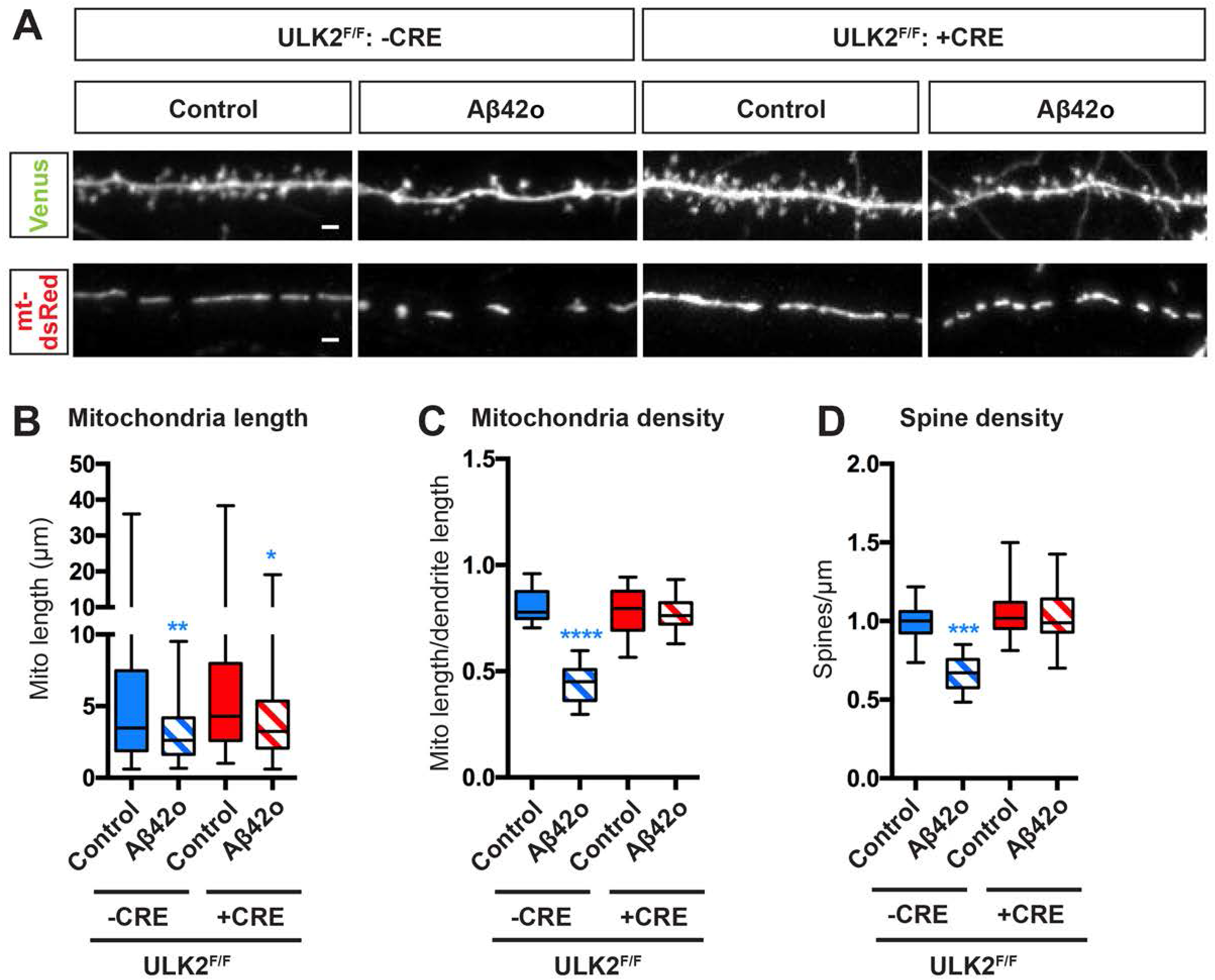
ULK2 acts in a concerted manner with MFF and leads to loss of mitochondrial biomass following MFF-dependent mitochondrial fragmentation. (**A**) Representative images of secondary dendritic segments of primary cortical neurons at 21DIV. ULK2^F/F^ embryos were *ex utero* electroporated at E15.5 with plasmids encoding with Venus and mitoDsRed as well as either scrambled Cre (control; -Cre) or Cre recombinase (+Cre). Neurons were treated at 21 DIV with either control or Aβ42o for 24 hours. Deletion of ULK2 in cortical PNs blocks Aβ42o-induced loss of dendritic spines and loss of mitochondrial biomass. (**B-D**) Quantification of dendritic mitochondrial length (B), dendritic mitochondrial density (C), and dendritic spine density (D). Analysis was done blind to the experimental conditions by manual counting using FIJI. Statistical analysis was performed using one-way ANOVA with Kruskal-Wallis followed by Dunn’s post-hoc test in (B-D). Data is represented by box plots displaying minimum to maximum values, with the box denoting 25^th^, 50^th^ (median) and 75^th^ percentile from 3 independent experiments. N_-CRE DMSOcontrol_ = 36 dendrites, 193 mitochondria; n_-CRE Aβ42o_ = 38 dendrites; 254 mitochondria; n_+CRE DMSOcontrol_ = 36 dendrites; 186 mitochondria; n_+CRE Aβ42o_ = 35 dendrites, 435 mitochondria. The test was considered significant when p<0.05 with the following criteria: * p<0.05; ** p<0.01; ***p<0.001; ****p<0.0001; ns, not significant. Scale bar for magnified dendritic segments= 2μm.

### AMPK phosphorylation of ULK2 is required for Aβ42o-induced dendritic mitophagy and synaptic loss

We next tested if AMPK-mediated phosphorylation of ULK2 is required for Aβ42o-dependent synaptotoxic effects in cortical PNs. Extensive studies in non-neuronal cells have established that the closest ortholog of ULK2, ULK1, is phosphorylated and thereby catalytically activated by AMPK and plays a key role in regulating mitophagy (Egan et al., 2011; Kim et al., 2011). Although the two orthologs seem to have redundant functions in non-neuronal cells, the above results uncovered that, in cortical PNs that only downregulation of ULK2, but not ULK1, completely blocks Aβ42-mediated synaptotoxicity (Fig. S6). Therefore, we performed a sequence homology analysis of candidate AMPK sites identified in ULK1 and compared them to ULK2. Two independent groups collectively reported six AMPK-mediated phosphorylation sites on ULK1 (Egan et al., 2011; Kim et al., 2011). Using primary sequence alignment between human and mouse ULK1/2, we have found that four of these ULK1 phosphorylation sites are conserved in ULK2 and are predicted as *bona fide* AMPK consensus phosphorylation sites (Fig.8A and Fig. S7; S309/T441/S528/S547). To verify that these sites are the four most prevalent AMPK target sites in ULK2, we mutated these four candidate S/T residues to alanine in ULK2 (ULK2-4SA) and over-expressed them in HEK293T cells. We then performed immunoprecipitation (IP) of Flag-tagged ULK2 WT and 4SA mutant from cells that are either treated with DMSO or AMPK direct agonist 991. Overexpression of constitutively active AMPK (CA-AMPK) in the presence of ULK2 WT or ULK2 4SA mutant conditions were used as a positive control. Immunoprecipitated Flag-ULK2 WT or Flag-ULK2 4SA were then immunoblotted with an antibody recognizing the optimal AMPK phospho-substrate motif. We observed basal levels of phosphorylation of ULK2 WT in untreated control which increases upon AMPK activation or when co-expressed with CA-AMPK (Fig. 8B). Importantly, ULK2 phosphorylation was completely abolished in the 4SA mutant (Fig. 8B), strongly suggesting that these four S/T sites are the main AMPK phosphorylation sites in ULK2.

**Figure 8.**
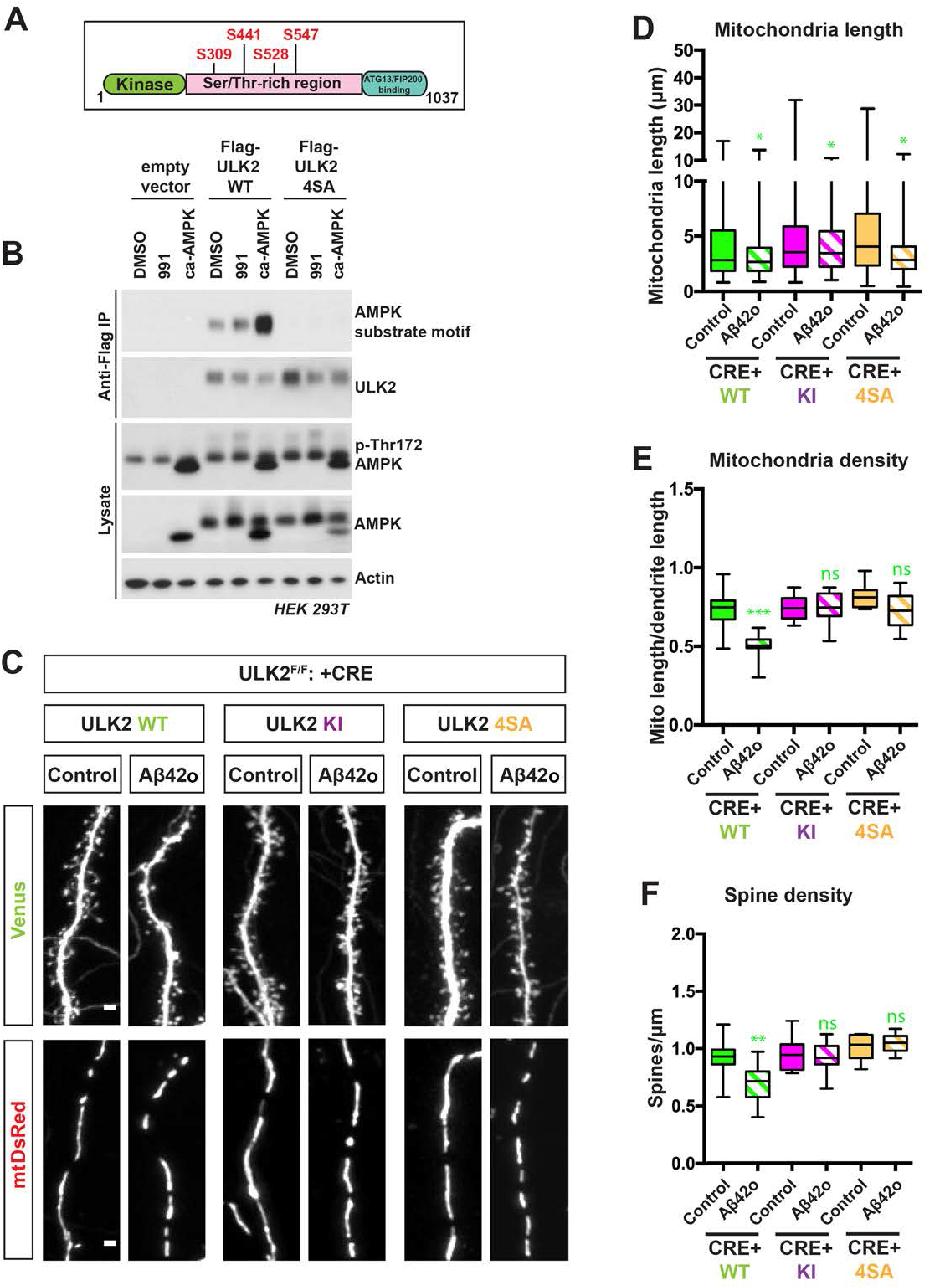
AMPK mediated phosphorylation of ULK2 is crucial for oligomeric Aβ42 induced synaptotoxicity and loss of mitochondrial biomass. (**A**) Schematic of mULK2 domain structure highlighting the four predicted AMPK-mediated phosphorylation sites (see Fig. S5B) conserved in ULK1 and ULK2. (**B**) Flag-mULK2 WT or mULK2 4SA (the four conserved phosphorylation sites shown in Fig. S5B are mutated to alanine) is over-expressed in HEK293T cells co-expressing either GST or GST-caAMPKα1(1-312) (constitutively active AMPK). Cells were treated with DMSO or 50μM compound 991 for 1hour. 5mg of whole cell lysate was immunoprecipitated with anti-Flag M2 Agarose and immunoblotted with AMPK substrate motif antibody and Flag M2 monoclonal antibody. 25μg of lysates were immunoblotted for p-Thr172 AMPK, total AMPK and actin. (**C**) Representative high magnification images of dendritic segments showing dendritic spines in the upper panel and mitochondria in the lower panel. ULK2^F/F^ embryos were *ex utero* electroporated at E15.5 with control vector (-Cre) or Cre recombinase encoding plasmid along with plasmids expressing Venus and mito-DsRed. The Cre-reconstituted (gene replacement) conditions were also co-electroporated with either mULK2 WT (Wild-type), mULK2 KI (K39I) (Kinase inactive) or mULK2 4SA (see Fig. S5). Neurons were treated at 20DIV with either control or Aβ42o for 24 hours. Abolishing ULK2 kinase-activity (ULK2 KI) or mutating AMPK-mediated phosphorylation sites on ULK2 (ULK2 4SA) can block Aβ42 oligomer induced synaptotoxicity and loss of mitochondrial biomass. (**D**) Quantification of dendritic mitochondrial length, and (**E**) dendritic mitochondrial density, and (**F**) spine density. Analysis was done blind to the experimental conditions by manual counting using FIJI. Statistical analysis was performed using one-way ANOVA with Kruskal Wallis followed by Dunn’s post-hoc test in (D-F). Data is represented by box plots displaying minimum to maximum values, with the box denoting 25^th^, 50^th^ (median) and 75^th^ percentile from 3 independent experiments. N_ULK2WT DMSOcontrol_ = 32 dendrites, 194 mitochondria; n_ULK2WT Aβ42o_ = 30 dendrites; 289 mitochondria; n_ULK2KI DMSOcontrol_ = 36 dendrites; 183 mitochondria; n_ULK2KI Aβ42o_ = 34 dendrites, 449 mitochondria; n_ULK24SA DMSOcontrol_ = 34 dendrites; 198 mitochondria; n_ULK24SA Aβ42o_ = 32 dendrites, 472 mitochondria. The test was considered significant when p<0.05 with the following criteria: * p<0.05; ** p<0.01; ***p<0.001; ****p<0.0001; ns, not significant. Scale bar for magnified dendritic segments= 2μm.

Finally, we tested if ULK2 phosphorylation by AMPK is required for the loss of mitochondrial biomass and the synaptic loss induced Aβ42o application. We performed gene replacement experiments taking advantage of ULK2^F/F^ primary cortical PNs co-electroporated with Cre (or delta Cre as negative control) along with ULK2-WT (positive control) or ULK2-4SA or kinase inactive ULK2 (ULK2-KI as negative control). We found that expression of ULK2-WT in ULK2-null cortical PNs enabled both loss of mitochondrial biomass and synaptic loss induced by Aβ42o, but the re-introduction of ULK2 KI or ULK2 4SA had no effect on mitochondrial biomass and did not prevent dendritic spine loss (Fig. 8C-F). These results confirm that AMPK mediated phosphorylation and activation of ULK2 mediate Aβ42o-dependent synaptic loss through its ability to induce mitophagy and loss of mitochondrial biomass.

### Phosphorylation of Tau on Serine 262 is required for Aβ42o-induced mitochondrial remodeling and synaptotoxicity

We and others previously shown that AMPK phosphorylates human Tau (hTau) on the KGxS motif present at Serine 262 embedded within the R1 microtubule binding domain (Mairet-Coello and Polleux, 2014; Thornton et al., 2011) and that expressing a form of hTau that cannot be phosphorylated at S262 (hTau-S262A) blocked Aβ42o-induced synaptic loss in cortical and hippocampal PNs (Mairet-Coello and Polleux, 2014). Since hTau phosphorylation has been involved in Drp1-mediated mitochondrial fission in multiple fly and mouse AD models (DuBoff et al., 2012; Manczak and Reddy, 2012), we tested specifically if AMPK-dependent hTauS262 phosphorylation could link Aβ42o-induced mitochondrial remodeling and synaptotoxicity. Indeed, expression of hTau-S262A (but not wildtype hTau) protected neurons from Aβ42o-induced mitochondrial remodeling and synaptotoxicity (Fig. 9A-D). This result strongly suggests that AMPK-dependent Tau phosphorylation on S262 participates to the MFF-ULK2-dependent mitochondrial remodeling and synaptic loss triggered by Aβ42o.

**Figure 9.**
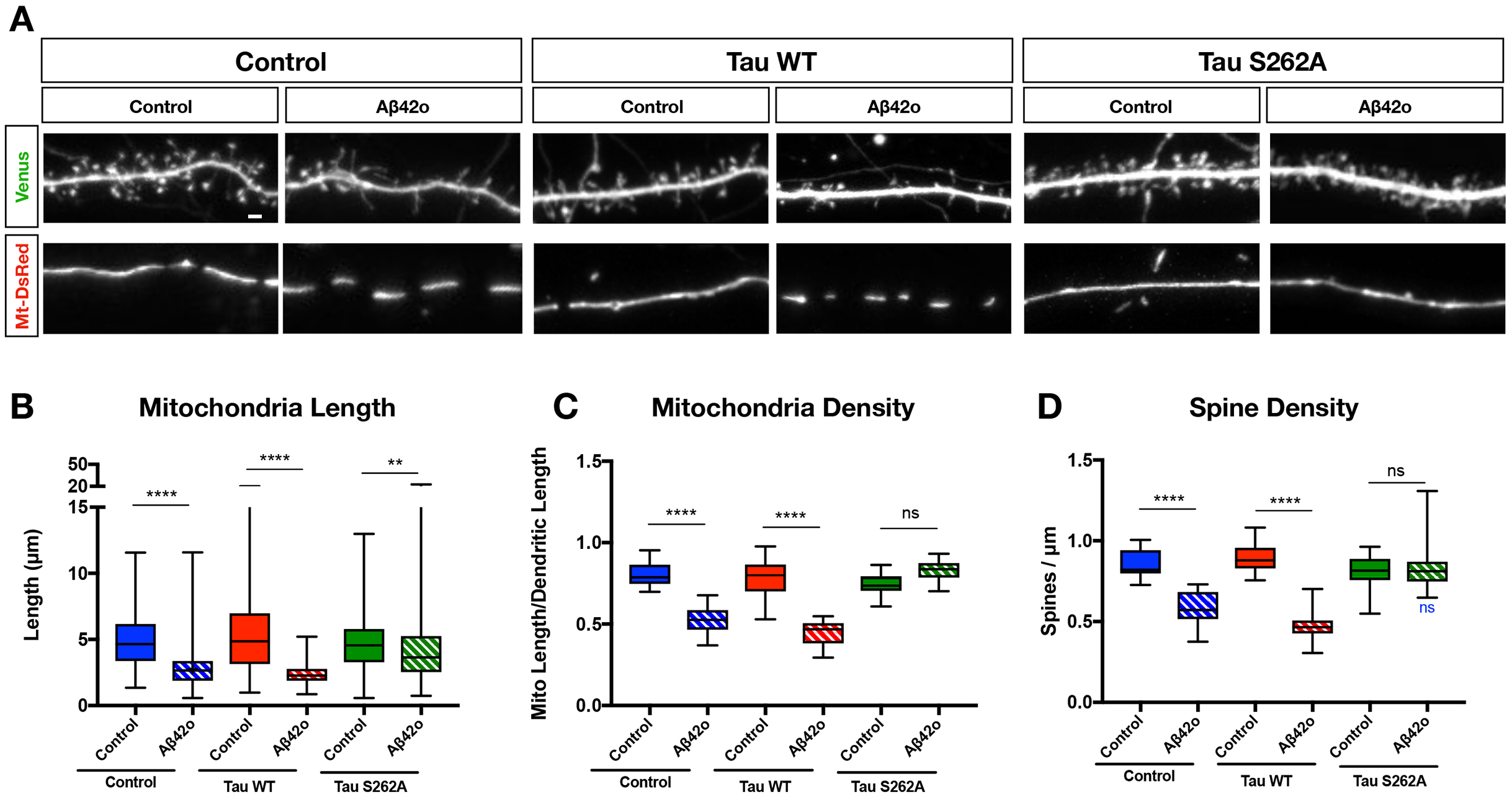
Tau phosphorylation at S262 is required for Aβ42o-induced dendritic mitochondrial remodeling. (**A**) Representative images of secondary dendritic segments of primary cortical PNs at 21 DIV. E15.5 mouse embryos were *ex utero* electroporated with plasmids encoding Venus, mitoDsRed only or in combination with WT hTau or a non-phosphorylatable mutant hTau-S262A. Neurons were treated at 20 DIV with either vehicle or Aβ42o at 1μM equivalent for 24 hours. Overexpression of hTau in the presence of Aβ42o decreases spine density, mitochondrial length, and mitochondrial density, while over expression of hTauS262A blocked oligomer-induced spine loss and a decrease in density, but not mitochondrial fragmentation. (**B**) Quantification of dendritic mitochondrial length, (**C**) dendritic mitochondrial density, and (**D**) dendritic spine density. All of the analyses were done blind to the experimental conditions, and was done by manual counting in FIJI. All of the statistical analyses were performed in Prism 6 (GraphPad Software). Data is represented by box plots displaying minimum to maximum values, with the box denoting 25th, 50th (median) and 75th percentile from 3-6 independent experiments. n_control_= 81 dendrites, 593 mitochondria; n_control Aβ42o_ = 105 dendrites, 645 mitochondria; n_hTau WT control_= 38 dendrites, 258 mitochondria; n_hTau WT Aβ42o_ = 79 dendrites, 725 mitochondria; n_hTauS262A control_ = 81 dendrites, 555 mitochondria; n_hTauS262A Aβ42o_ = 89 dendrites, 756 mitochondria. Statistical analyses were performed using One-way ANOVA followed by Kruskal-Wallis Multiple Comparisons in (B-D). The test was considered significant when p < 0.05 with the following criteria: * p < 0.05; ** p < 0.01; *** p < 0.001; **** p < 0.0001; ns, not significant.

## Discussion

Loss of excitatory glutamatergic synapses in cortical and hippocampal PNs has been reported as an early event in AD progression, primarily driven by soluble Aβ42o and Tau hyperphosphorylation (Hsieh et al., 2006; Ittner et al., 2010; Lacor et al., 2007; Mucke et al., 2000a; Mucke et al., 2000b; Roberson et al., 2011; Roberson et al., 2007; Shankar et al., 2007; Sheng et al., 2012). This loss of connectivity is thought to be responsible for the early cognitive defects characterizing AD patients including defective learning and memory (Sheng et al., 2012; Terry et al., 1991). Recently, mitochondrial remodeling and dysfunction in PNs have emerged as a prominent cellular phenotype in AD (Cho et al., 2009; Fang et al., 2019; Wang et al., 2009a; Wang et al., 2009b; Zhang et al., 2016). However, whether this structural remodeling of dendritic mitochondria is causally linked to excitatory synaptic loss was unknown, in part because the molecular mechanism mediating Aβ42o-dependent mitochondrial remodeling are currently unknown.

AMPK is best-characterized as a metabolic sensor in non-neuronal cells (Herzig and Shaw, 2018). Importantly, in AD patients, catalytically active AMPK is abnormally accumulated in the cytoplasm of PNs of CA1, entorhinal cortex, and neocortex (Vingtdeux et al., 2011). AMPK over-activation has been consistently reported in AD patient brain samples (Ma et al., 2014; Vingtdeux et al., 2011), in various AD mouse models (Ma et al., 2014; Mairet-Coello et al., 2013; Son et al., 2012), as well as *in vitro* upon acute treatment of Aβ42o (Ma et al., 2014; Mairet-Coello et al., 2013; Thornton et al., 2011; Yoon et al., 2012). The CAMKK2-AMPK signaling pathway is unique in its involvement in multiple phenotypes observed during the progression of AD. The present study and previous reports collectively implicate this signaling pathway as the first to unify multiple phenotypes reported in the brain of AD patients and in AD mouse models, including disruption of Ca^2+^ homeostasis, APP processing, hTau phosphorylation, mitochondrial remodeling, increased autophagy, and synaptic loss (Marinangeli et al., 2018; Nixon, 2005, 2013; Sheng et al., 2012). There is growing evidence that one of the earliest event upon Aβ42o application is disrupted cytoplasmic Ca^2+^ homeostasis that drives PNs hyperexcitability (see below) (Arbel-Ornath et al., 2017; Busche et al., 2012; Grienberger et al., 2012; Kuchibhotla et al., 2008; Lerdkrai et al., 2018; Palop and Mucke, 2010). This increased Ca^2+^ accumulation precedes spine morphological changes (Arbel-Ornath et al., 2017), strongly suggesting that Ca^2+^-dependent signaling pathways, including the CAMKK2-AMPK pathway, occur early in the disease progression and plays a critical role in reduction of synaptic plasticity and synaptic maintenance (Hsieh et al., 2006; Ma et al., 2014; Malinow, 2012). Interestingly, Aβ42o-induced AMPK over-activation has been found to induce a vicious cycle of increased Aβ42o production via activation of JNK3-dependent APP processing (Yoon et al., 2012). Furthermore, AMPK links the two key players of AD: Aβ42o and hTau. Catalytically active AMPK has been shown to co-localize with phosphorylated-S262 hTau in brains of AD patients (Vingtdeux et al., 2011), and previous work highlights that hyperactivation of AMPK by Aβ42o and /or other AMPK-like kinases such as NUAK1 leads to Tau phosphorylation on two of its serine residues (S262 and S356; (Lasagna-Reeves et al., 2016; Mairet-Coello et al., 2013)) embeded in the R1 and R4 microtubule binding domains respectively (Buee et al., 2000). These results identified for the first time the CAMKK2-AMPK signaling pathway as a critical link between Aβ42o to Tau phosphorylation in the induction of synaptic loss (Mairet-Coello et al., 2013; Mairet-Coello and Polleux, 2014; Thornton et al., 2011).

In this paper, we identified two new critical effectors of AMPK mediating the effects of Aβ42o on synaptic loss. We show that AMPK over-activation by Aβ42o coordinates multiple aspects of mitochondrial remodeling by co-regulating mitochondrial fission and mitophagy through its activation of MFF and ULK2, respectively. For the first time, we demonstrate that Aβ42o mediates loss of excitatory synapses through MFF-ULK2-dependent mitochondrial fission and mitophagy. We have identified that Aβ42o-mediated AMPK over-activation also signal through phosphorylation of Tau on S262 in parallel to MFF-ULK2 phosphorylation/activation and lead to dendritic mitochondrial remodeling and synapse loss. Further work will be required to understand if Aβ42o-induced AMPK-Tau signaling plays a direct role in mitochondrial fission and/or mitophagy machinery or plays an indirect role through its postsynaptic signaling function (Ittner and Gotz, 2011; Ittner et al., 2010) and/or its ability to regulate microtubule dynamics (Qu et al., 2017) thereby indirectly influence mitochondrial remodeling.

Lastly, what makes this molecular mechanism attractive for development of new therapeutic approaches is that it is a true bonafide stress-response signaling pathway (Mairet-Coello and Polleux, 2014) i.e. these AMPK-dependent pathways are dispensable for normal neuronal development and/or synaptic maintenance in adult cortical and hippocampal neurons (Mairet-Coello et al., 2013; Mairet-Coello and Polleux, 2014; Williams et al., 2011). Genetic manipulations of AMPK or its downstream effectors do not have any identified phenotypic effects on neurons under basal conditions, but only upon induction of metabolic stress or Aβ42o application in neurons.

Mitochondrial remodeling and dysfunction are converging mechanisms of disease pathogenesis shared by multiple neurodegenerative diseases (ND). This is in part explained by the disruptions of mitochondrial homeostasis, motility, and dynamics observed in various ND as well as genetic evidence suggesting a strong association between genes involved in mitochondrial function and various adult on-set ND syndromes (Itoh et al., 2013; Schon and Przedborski, 2011). In AD specifically, altered mitochondrial dynamics have been implicated by the changes of mitochondrial fission and fusion protein expression levels in AD patient brains (Wang et al., 2009a) and various AD mouse models (Wang et al., 2008), as well as altered posttranslational modification of Drp1 thought to disrupt mitochondrial dynamics (Cho et al., 2009; Wang et al., 2009a). In line with our results, Wang, et al., (2009) also observe that in the presence of Aβ42o, dendritic mitochondria decrease in length and density, which is correlated with reduced spine density. Interestingly, a recent study suggested a significant degree of mitochondrial structural remodeling in various AD mouse models and AD patients in the CA1 of the hippocampal region using 3D EM reconstructions (Zhang et al., 2016). Our present study further corroborates these observations and provides a molecular mechanism mediating Aβ42o-dependent mitochondrial remodeling and synaptic loss during early stages of AD.

A second cellular defect thought to contribute to several neurodegenerative diseases including AD is a disruption of autophagy-lysosomal pathway (Nixon, 2013). Although the cellular and molecular mechanisms of autophagy have been examined using non-neuronal cells (Mizushima, 2007), recent results demonstrated unique, compartment-specific regulation of autophagy in neurons. Local biogenesis of autophagosomes has been observed in distal axons and traffic retrogradely towards the soma in order to fuse with lysosomes, whereas autophagosomes in the soma are more stationary (Maday and Holzbaur, 2016). However, the dynamics of autophagy in dendrites remains largely unexplored. Much of what we know on neuronal autophagy has been primarily focused on axonal and presynaptic autophagy dynamics, where it mediates efficient synaptic transmission by regulating the turnover of synaptic proteins (Uytterhoeven et al., 2015). Interestingly, autophagosome (AP) formation occurs in distal axons and can traffic retrogradely to fuse with lysosomes that are localized closer to the soma (Maday and Holzbaur, 2016). Autophagy has received significant attention in AD research, in part due to the abnormal accumulation of AP and autolysosomes (AL) in dystrophic neurites of AD patients (Nixon, 2005). Moreover, familial mutations and polymorphisms associated with AD link to dysfunction of the autophagic-lysosomal pathway. Mutations of the Presenillin (PS) 1 and 2 proteins have been shown to disrupt lysosomal acidification and autophagy (Lee et al., 2015). AD associated genes such as PICALM, BIN1, RAB11, and VPS34 are involved in various initiation, sorting, and trafficking processes of autophagy (Ando et al., 2016; Miyagawa et al., 2016; Morel et al., 2013; Udayar et al., 2013). However, the majority of the cellular phenotypes involving AP and AL buildup in neurons have been described at late stages of AD progression when neurodegeneration involving actual neuronal cell loss occurs (Nixon, 2005). More recent results suggested that, in AD mouse models, autophagic flux is disrupted (Son et al., 2012) and expression level of mitophagy genes are elevated during the early stages of AD (Sorrentino et al., 2017). Our results demonstrate that Aβ42o triggers an AMPK-ULK2 dependent increase in mitophagy resulting in a significant local degradation of dendritic mitochondria and most interestingly, that this cell biological step is causally linked with synaptic loss since we rescue dendritic spine loss induced by Aβ42o in cortical PNs where ULK2 phosphorylation by AMPK is disabled.

Our *in vivo* analysis also reveals a striking and previously unknown degree of compartmentalization of mitochondria morphology in dendrites of CA1 PNs: in basal and apical oblique dendrites, mitochondria are small and punctate whereas the distal apical dendrites contain elongated and fused mitochondria (Fig. S1). The transition between these two types of mitochondrial morphology is sharp and actually corresponds to the transition between two hippocampal layers, stratum radiatum (sr) and stratum lacunosum moleculare (slm) where the earliest excitatory synaptic loss is observed in this mouse model. This is particularly relevant because the synapse made between medial entorhinal cortical inputs and the apical tufts of CA1 PNs in stratum lacunosum moleculare has been described as one of the central synapse defective in late onset forms of AD (Small et al., 2011). Future investigations will need to determine why this synapse is particularly vulnerable during early stages of AD progression.

As discussed above, disruption of Ca^2+^ homeostasis has been observed in various AD mouse models (Busche et al., 2012; Grienberger et al., 2012; Lerdkrai et al., 2018), in wildtype PNs exposed to acute treatment of oligomeric Aβ42 *in vivo* (Arbel-Ornath et al., 2017), and neuronal culture exposed to Aβ42o *in vitro* (Demuro et al., 2005; Guo et al., 1999). Recent results suggested that disruption of Ca^2+^ dynamics is largely due to Aβ42o, as immune-depletion of Aβ42o can prevent this phenotype (Arbel-Ornath et al., 2017) and increase the population of hyperactive neurons (Busche et al., 2012; Grienberger et al., 2012). Possible explanations of this increased cytoplasmic Ca^2+^ accumulation include increased mobilization of Ca^2+^ from extracellular space through NMDAR and VGCC, increased leakage of Ca^2+^ from intracellular Ca^2+^ storing organelles such as the ER, and loss of Ca^2+^ buffering capacity by the neurons. Currently, there is evidence supporting that Ca^2+^ is derived from the extracellular influx of Ca^2+^ via NMDAR (Arbel-Ornath et al., 2017; Sheng et al., 2012) as well as the Ca^2+^ released from the ER via the IP_3_R and RyR (Chan et al., 2000; Stutzmann et al., 2004). Regardless of the Ca^2+^ source, PNs ultimately experience intracellular Ca^2+^ increase that can activate Ca^2+^-dependent signaling pathway, including our CAMKK2-AMPK pathway. There may be a complex positive feed-forward loop where Aβ42o-dependent, fast NMDAR-dependent Ca^2+^ influx (Arbel-Ornath et al., 2017) over-activates the CAMKK2-AMPK pathway (Mairet-Coello et al., 2013), resulting in reduction of mitochondrial biomass, i.e. mitochondrial matrix volume, and therefore could lead to reduced Ca^2+^ buffering capacity, further increasing cytoplasmic Ca^2+^ accumulation and CAMKK2 over-activation. Future work will need to address if the striking but spatially restricted degree of Aβ42o-dependent reduction in dendritic mitochondrial biomass and the reduction in spine density identified in the present study in the apical tuft dendrites of CA1 PNs in the J20 AD mouse model contributes to degradation of the spatial tuning properties of these neurons at early stages of the disease progression.

## Acknowledgments

We thank members of the Polleux lab for feedback and discussion, as well as Karen Duff, Ulrich Hengst, Carol Troy for critically evaluating the manuscript. We thank Benoit Viollet (INSERM, France) for providing the AMPK double conditional mouse lines. We thank Fan Wang (Duke University) for providing ULK1 and ULK2 shRNA constructs. We thank Qiaolian Lu and Miyako Hirabayashi for excellent management of our mouse colony. This work was supported by grants from the NIH (NS089456) (FP), a F31 Award from NIH (AL), a K99 award (NS091526) (TLL), the Henry and Marilyn Taub Foundation (AS), the Ludwig Foundation (FP, AS) and an award from the Fondation Roger De Sproelberch (FP).

## Material and methods

### Animals

Mice were used according to protocols approved by the Institutional Animal Care and Use Committee (IACUC) at Columbia University and in accordance with National Institutes of Health guidelines. Time-pregnant CD1 females were purchased from Charles Rivers. 129/SvJ, C57Bl/6J nontransgenic mice and hemizygous transgenic mice from line J20 (hereafter referred as J20) (The Jackson Laboratory stock #006293) were maintained in a 12-hour light/dark cycle. J20 mice express human APP carrying the Swedish and Indiana mutations under a PDGFβ promoter (Mucke et al., 2000a; Palop et al., 2007). AMPKα1^F/F^α2^F/F^ double conditional knockout mice were a generous gift from Dr. Benoit Viollet (INSERM, Institut Cochin, Paris-France). Ulk2^F/F^ mice were obtained from The Jackson Laboratory stock #017977 (Cheong et al., 2011). Timed-pregnant females were obtained by overnight breeding with males of the same strain. Noon the day after the breeding was considered as E0.5.

### Synthetic Aβ42 oligomers preparation

Aβ42 (rPeptide or Creative Peptide) was processed to generate Aβ42 oligomers as described previously (Mairet-Coello et al., 2013). Briefly, lyophilized Aβ42 peptide was dissolved in hexafluoro-2-propanol (HFIP; Sigma-Aldrich) for 2 hours to allow monomerization. HFIP was removed by speed vacuum, and the monomers were stored in −80°C. Aβ42 monomers were dissolved in anhydrous dimethyl sulfoxide (DMSO) to make a 5mM solution, then added to cold phenol red-free F12 medium (Life Technology) to make a 100uM solution. The solution was incubated at 4°C for two days and centrifuged at 12,000 × g for 10 min at 4°C in order to discard the fibrils. The supernatant containing the 4°C oligomers were assayed for protein content using the BCA kit (Thermo Fisher Scientific). For control experiments, vehicle treatment corresponded to the same volume of DMSO and F12 media used for Aβ42o treatment and for Figure S2, a peptide corresponding to the inverted sequence of Aβ42 (INV42; rPeptide and Creative Peptide) was processed as for Aβ42 oligomerization. For western blotting of Aβ42 oligomers, 16.5% Tris-Tricine SDS-PAGE was performed. Synthetic Aβ42 oligomers were prepared in 2X Tricine sample buffer (BIORAD) without reducing agent and resolved by SDS-PAGE. The separated proteins were transferred onto Immobilin-FL PVDF membrane (EMD Millipore), blocked for 1 hour with Odyssey blocking buffer (LICOR) and probed with 6E10 monoclonal antibody (Covance) overnight at 4°C. Secondary antibody incubation was performed for 1 hour at room temperature with IRDye 680RD goat anti-mouse secondary (LICOR) and the blot was visualized using the Odyssey Imaging system. To estimate the relative concentration of synthetic Aβ42 peptide monomer versus oligomers (dimers and trimers), we performed quantitative western blotting (see Fig. S2E) and measured the ratio of optical density measured for the monomeric peptide versus the dimer/trimer bands. Using this approach, we estimated that a 1μM concentration of oligomerized peptide, contains approximately 300-450nM of effective dimers/trimers.

### Primary cortical culture and *ex utero* electroporation

Cortices from E15.5 mouse embryos were dissected followed by dissociation in complete Hank’s balanced salt solution (cHBSS) containing papain (Worthington) and DNase I (100ug/mL, Sigma) for 15 minutes at 37°C, washed three times, and manually triturated in DNase I (100ug/mL) containing neurobasal medium (Life Technology) supplemented with B27 (1x, Thermo Fischer Scientific,), FBS (2.5%, Gibson) N2 (1x, Thermo Fischer Scientific), glutaMAX (2mM, Gibco). Cells were plated at 10.0 × 10^4^ cells per 35 mm glass bottom dish (Mattek) that has been coated with poly-D-lysine (1mg/mL, Sigma) over night. One-third of the medium was changed every 5 days thereafter with non-FBS containing medium and maintained for 20-25 days in 5% CO_2_ incubator at 37°C. Ex utero electroporation was performed as previously published (Courchet et al., 2013). See the Supplemental Experimental Procedures for details and constructs. Plasmids used for *ex utero* electroporation where all in pCAG vector backbone (Guerrier et al., 2009) expressing the following cDNAs: LAMP1-mEmerald (Lewis et al., 2016), mito-DsRed and Venus (Courchet et al., 2013), mRFP-LC3 (originally from https://www.addgene.org/21075/ and subcloned into pCAG).

### *In utero* hippocampal electroporation

*In utero* electroporation targeting the hippocampus was performed as previously described (Hand and Polleux, 2011; Mairet-Coello et al., 2013) with slight modifications as described in (Szczurkowska et al., 2016) to target the embryonic hippocampus at E15.5. See Supplemental Experimental Procedures for more details.

### Imaging

Imaging on dissociated neurons was performed between 20-25 DIV in 1,024 x 1,024 resolution with a Nikon Ti-E microscope equipped with A1R laser-scanning confocal microscope using the Nikon software NIS-Elements (Nikon, Melville, NY, USA). We used the following objective lenses (Nikon): 10x PlanApo; NA 0.45 (for images of hippocampal slices), 60X Apo TIRF; NA 1.49 (for analysis of spine density and mitochondrial morphology in cultured neurons), 100x H-TIRF; and NA 1.49 (for analysis of spine densities and mitochondrial morphology in brain slices). Dendritic spine density was quantified on secondary or tertiary dendritic branches that were proximal to the cell body, on z projections for cultured neurons and in the depth of the z stack for slices, using FiJi software (ImageJ; NIH). See Supplemental Experimental Procedures for more details.

### hESC culture

H9 (WA09; WiCell) and its genome edited derivatives were maintained on StemFlex (Life) and Cultrex substrate (Biotechne), and routinely split ∼ twice a week with ReLeSR (Stem Cell Technologies) in the presence of ROCKi (Y-27632; Selleckchem). H9 is a commercially available hESC line on the NIH Registry (# 0062) and Dr. Sproul has approval to genome edit and differentiate this hESC line by the Columbia University Human Embryonic and Human Embryonic Stem Cell Research Committee.

### APP^Swe^ knockin hESC line generation

The APP^Swe^ mutation (KM670/671NL) was knocked into both alleles of H9 using CRISPR/Cas9, bi-allelic knockin of APP^Swe^ has been demonstrated to have increased A*β* production relatively to monoallelic knockin in a control iPSC line (Paquet et al., 2016). In brief, a sgRNA targeting Exon16 of *APP* was designed and subcloned into the MLM3636 vector, a gift from Keith Joung (Addgene plasmid # 43860; http://www.addgene.com/43860). A ssDNA HDR template (IDT) was designed in which the APP^Swe^ mutation disrupted the PAM and introduced a *de novo* Xba1 restriction site. The sgRNA, Cas9-GFP (a gift from Kiran Musunuru (Addgene plasmid # 44719; http://www.addgene.org/44719/), and ssDNA HDR template were electroporated (Lonza nucleofector) into feeder-free H9 hESCs, followed by cell sorting on GFP signal and plating at low density on MEFs (MTI-GlobalStem). Individual clones were manually picked into a 96 well format, subsequently split into duplicate plates, one of which were used to generate sgDNA as had been done previously (Paquet et al., 2016). For each clone, exon 16 of *APP* was amplified and initially screened by restriction digest with Xba1 (NEB). Sanger sequencing was used to confirm the mutation, and successful knockin clones were expanded and banked. Potential off-target effects of CRISPR/Cas9 cleavage were analyzed by Sanger sequencing of the top 5 predicted off-target genomic locations [https://mit.crispr.edu], which demonstrated a lack of indels for multiple clones. One APP^Swe/^APP^Swe^ knockin line was analyzed and demonstrated to have a normal karyotype (G-banding, Columbia University Clinical Cytology Laboratory), and was used in the present work (Cl. 160).

### Transdifferentiation of hESCs into cortical-like pyramidal neurons

pLV-TetO-hNGN2-eGFP-Puro was generously provided by Kristen Brennand (Ho et al., 2016) and FUdeltaGW-rtTA was a gift from Konrad Hochedlinger (Addgene plasmid # 19780; http://www.addgene.org/19780/) (Maherali et al., 2008). Concentrated lentiviruses for these two plasmids were made using Lenti-X concentrator (Takara) as has been done previously (Sun et al., 2019). Permanent doxycycline inducible Ngn2-eGFP lines were generated by reverse infecting H9 and H9:APP^Swe/Swe^ hESC lines with NGN2-eGFP and rtTA and subsequent selection by puromycin.

For transdifferentiation of these two lines of hESCs into cortical neurons (iNs; (Zhang et al., 2013)), H9 and H9:APP^Swe/Swe^ lines were plated at ∼165,000 cells per 12 well-well in N2/B27 medium (Topol et al., 2015) on PEI (0.1% Sigma)/laminin (10 ug/mL, Biotechne) coated plates, supplemented with 1 ug/mL doxycycline (Sigma),10 ng/mL BDNF and NT3 (Biotechne) and 1 ug/mL laminin (Biotechne). BDNF, NT3 and laminin were added at similar levels for all subsequent feeds for the duration of the experiment. One day post plating cells were treated with puromycin (1ug/mL, Fisher). The following day cells were treated with 2 uM AraC (Biotechne) for 4 days, and 100 ng/mL on subsequent feeds. One week post infection cells were transitioned into BrainPhys (Stem Cell Technologies). Two weeks post infection, 10% mouse astrocyte conditioned media (ACM; ScienCell) was also added to the medium. Transfection of mitoDsRed2 was done at day 21-23 post infection using Lipofectamine 3000 (Life), and cells were fixed and analyzed 4 days later. A full media change (1/2 saved conditioned media) was performed ∼ 6h post transfection. Note that iNs had poor survival from Lipofectamine transfection without either ACM or primary mouse glia present.

### Cell Lysis and Immunoprecipitation

Neuron and differentiated hESC cultures were washed with cold PBS and lysed in Triton Lysis Buffer: 20mM HEPES (pH 7.5), 150mM NaCl, 1 mM EDTA, 1 mM EGTA, 1% Triton X-100, 0.25M Sucrose, Complete™protease inhibitor (Roche), and phosphoSTOP (Roche). Lysates were incubated at 4°C for 15 minutes and cleared at 15,000 × g for 15 minutes at 4°C. Total protein was normalized using Bio-rad Protein Assay Dye (Bio-rad).

For western blotting of phospho-T172 AMPK and total AMPK, equal amounts of lysates were loaded on a Mini-Protean TGX (4-20%) SDS-PAGE (Bio-rad). The separated proteins were transferred onto polyvinylidene difluoride membrane (PVDF, Bio-Rad), blocked for 1 hour with blocking buffer containing 5% fat-free dry milk in TBS-T. Membranes were then incubated overnight at 4°C with different primary antibodies diluted in the same blocking buffer. Incubations with HRP conjugated secondary antibodies were performed for 1 hour at room temperature, and visualization was performed by chemiluminescence. For phospho-MFF and MFF immunodetection, equal amounts of lysates (80μg) were immunoprecipitated with MFF or HA (as negative control for specific binding) to enhance MFF proteins. SureBeads^TM^ Protein A Magnetic Beads were washed three times with lysis buffer and added to equilibrated lysates for 1 hour at 4°C. The beads were washed 2 times with lysis buffer, two times with lysis buffer without Triton X-100, and then eluted by boiling in SDS lysis buffer for 5 minutes at 95°C and resolved by SDS-PAGE (Bio-rad) gel. The separated proteins were transferred onto polyvinylidene difluoride membrane (PVDF, Bio-Rad), blocked for 1 hour with blocking buffer containing 5% BSA in Tris-buffered saline solution. Membranes were incubated overnight at 4°C with either phospho or total MFF primary antibodies diluted in the same blocking buffer. Clean-Blot^TM^ IP Detection Kit (Thermo Fisher Scientific) was used to detect the bands of interest without the detection of denatured IgG as MFF protein size is close to both heavy and light chain. After overnight primary antibody incubation, membranes were incubated with Dilute Clean-Blot IP Detection Reagent at 1:100 for 1 hour at room temperature and visualization was performed by enhanced chemiluminescence.

For phospho-ULK2 experiments, 293T cells were transfected using Lipofectamine 2000 (Life Technologies) according to manufacturer’s instructions with pcDNA3 plasmids encoding flag tagged WT ULK2, flag tagged K39I ULK2 (ULK2 KI) or flag tagged S309A,T441A,S528A,S547A ULK2 (SA4). Where indicated, cells were co-transfected with a constitutively active truncated version of AMPK*α*1 (caAMPK).

Transfected cell lysates were harvested 40-48 hours post transfection. 1 hour after changing the media with fresh media containing vehicle or 50µM compound 991 (Glixx laboratories, Inc.), cells were washed with cold PBS and lysed in lysis buffer: 20 mM Tris pH 7.5, 150 mM NaCl, 1 mM EDTA, 1mM EGTA, 1% Triton X-100, 2.5 mM pyrophosphate, 50 mM NaF, 5 mM β-glycero-phosphate, 50 nM calyculin A, 1 mM Na3VO4, and protease inhibitors (Roche). Lysates were incubated at 4°C for 15 minutes and cleared at 16,000g for 15 minutes at 4°C. Total protein was normalized using BCA protein kit (Pierce) and lysates were resolved on SDS-PAGE gel. Immunoprecipitations were performed by adding magnetic M2 FLAG beads (Sigma) to equilibrated lysates for 2 hours at 4°C. The beads were washed three times with lysis buffer and then eluted by boiling in SDS lysis buffer for 5 minutes and resolved by SDS-PAGE.

Antibodies used in the paper are the following: anti-phospho-T172-AMPKα (40H9, 1:1000, Cell Signaling); AMPKα1/2 (1:1000, Cell Signaling); MFF (1:1000, Proteintech); phospho-S146/172-MFF (1:1000, Cell Signaling); pAMPK motif (CST, #5759S); total AMPK*α* (CST, #2532), Flag M2 (Sigma, F7425), pAMPK (CST, #2535), actin (Sigma, A5441). HRP-coupled secondary antibodies to mouse (AP124P) or rabbit (AP132P) were from Millipore. Li-Cor Fluorescence-coupled secondary antibodies to mouse or rabbit were from Li-Cor Biosciences.

### Statistics

Statistical analyses were performed with Prism 6 (GraphPad Software). The statistical test applied for data analysis is indicated in the corresponding figure legend. Experimental groups where all distributions were Gaussian/normal were assessed using the unpaired t-test for two-population comparison, or one-way ANOVA with Dunnett’s post hoc test for multiple comparisons. Nonparametric tests including the Mann-Whitney U test for two-population comparison and kruskal-Wallis with Dunn’s posttest for multiple comparisons were applied when distributions were not Gaussian. Unless otherwise noted, data are expressed as mean ± SEM. For dendritic spine and mitochondrial morphology analysis, all data were obtained from at least three independent experiments or at least three individual mice. The test was considered significant when p<0.05. For all analyses, the following apply: * p<0.05; ** p<0.01; *** p<0.001; **** p<0.0001; ns, not significant with p > 0.05.

## Supplemental Figures

**Figure S1.**
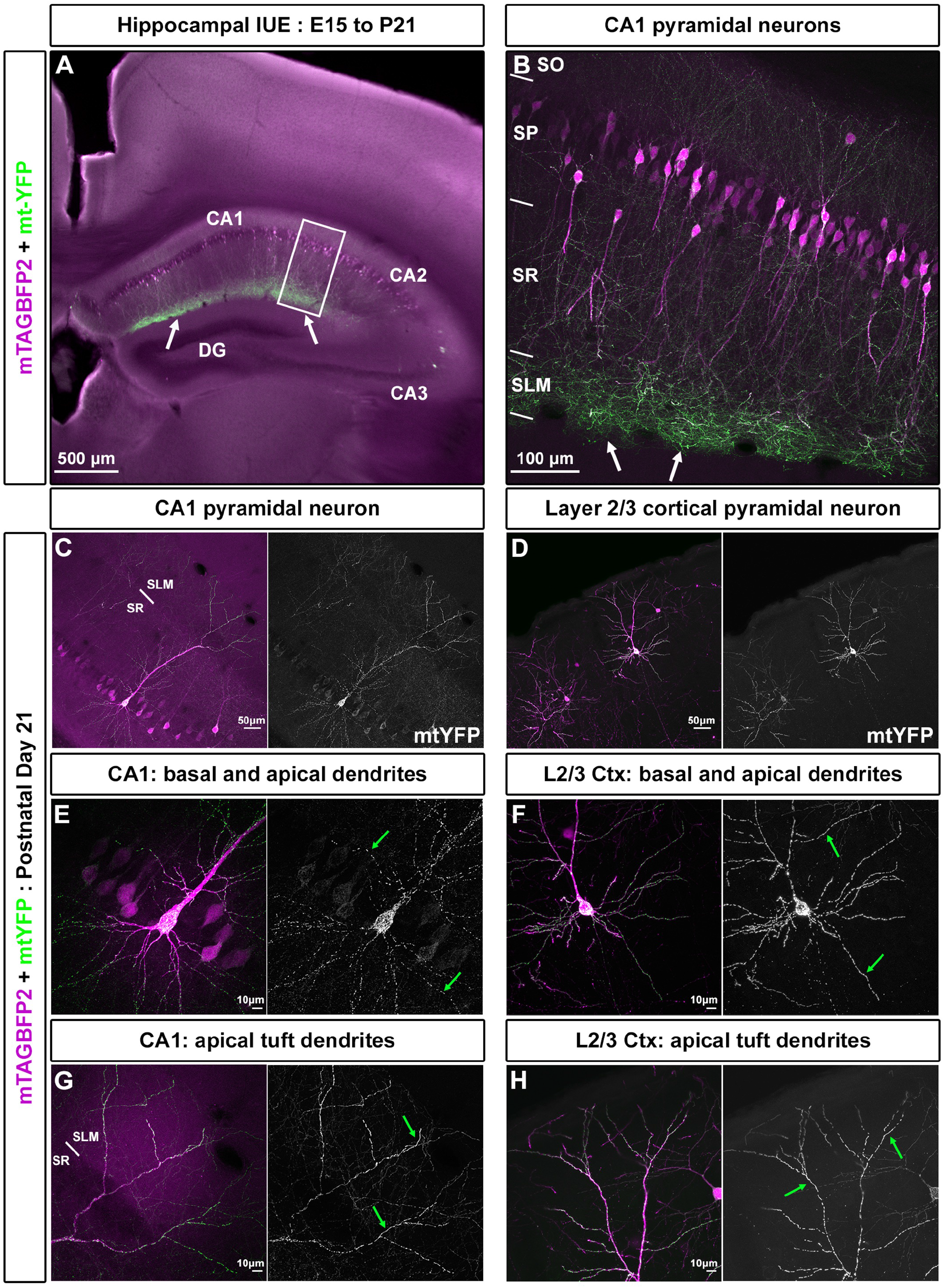
Hippocampal CA1 pyramidal neurons are characterized by highly compartmentalized dendritic mitochondria morphology compared to layer 2/3 cortical pyramidal neurons. ***Related to Figure 1*** (A) Low magnification image of hippocampal CA1 pyramidal neurons in utero electroporated at E15.5 with pCAG-mTagBFP2 (purple) and pCAG-mito-YFP (green) imaged at P21. (B) High magnification image of panel A highlighting that the distal apical dendrites in the SLM layer contain high density of mitochondria compared to the basal and oblique dendrites (in so and sr, respectively). (C-D) Low magnification of a single CA1 pyramidal neuron and a single Layer 2/3 cortical pyramidal neuron. (E) High magnification of CA1 PN shown in C highlighting that mitochondria in the basal and oblique dendrites are small and fragmented (arrowheads). (F) High magnification of cortical L2/3 PN shown in D, highlighting that mitochondria morphology is homogenously elongated (arrow) in basal and proximal apical dendrites. (G) High magnification of CA1 PN shown in C highlighting that mitochondria morphology in the distal apical tuft dendrites (slm) are elongated unlike those found in basal and oblique dendrites of the same cell (see E). (H) High magnification of cortical L2/3 PN highlighting that mitochondria morphology is homogenously elongated in distal apical dendrites similar to those found in both basal and oblique dendrites in the same neuron (see F). Abbreviations: slm, stratum lacunosum moleculare; sr, stratum radiatum; so, stratum oriens; sp, stratum pyramidalis.

**Figure S2.**
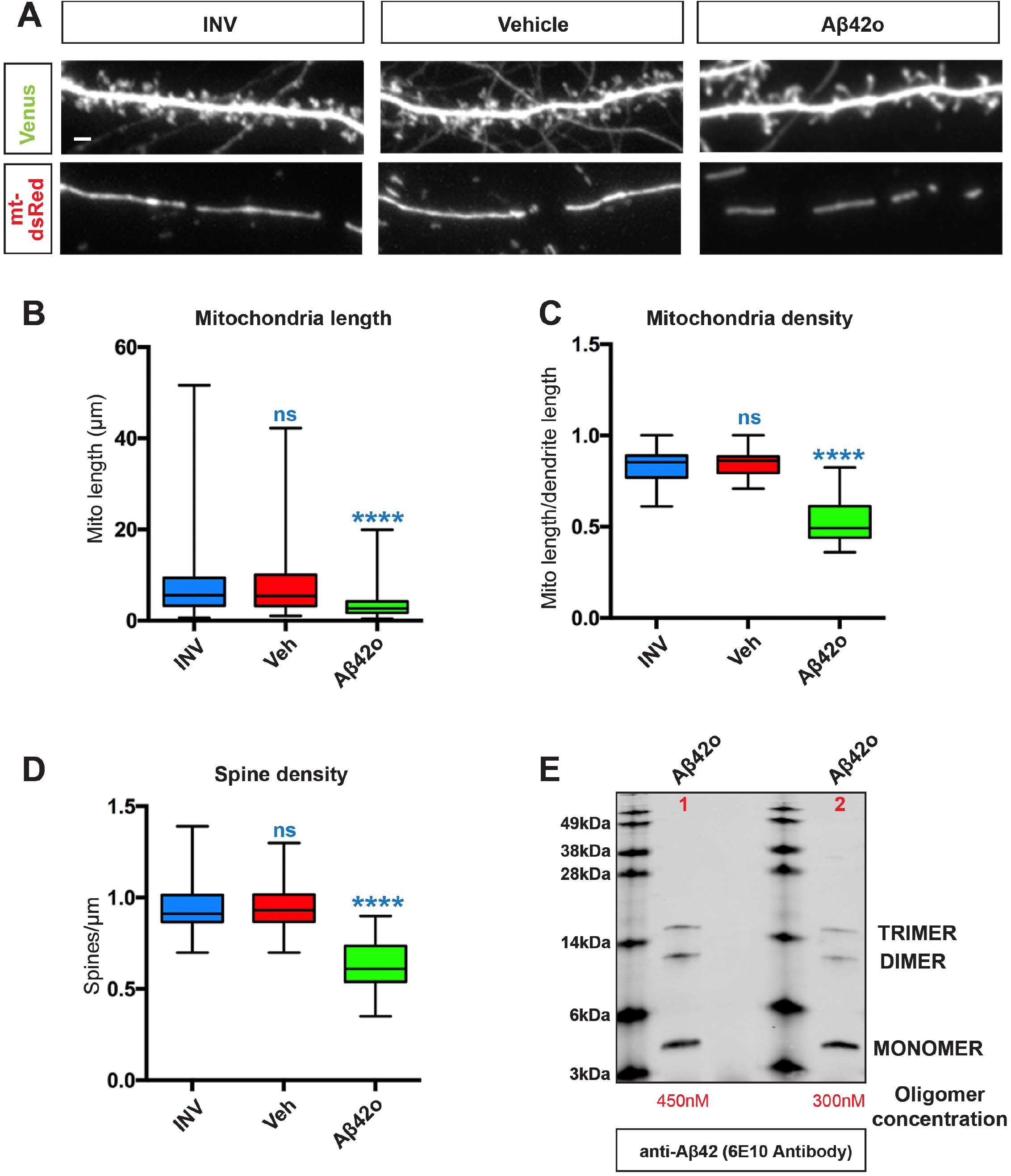
Treatment of inverse oligomeric Aβ42 does not influence mitochondrial morphology or spine density. ***Related to Figure 2*** (**A**) Representative high magnification images of dendritic segments showing dendritic spines in the upper panel and mitochondria in the lower panel. Treatment of inverse Aβ42o does not affect spine density or mitochondrial morphology and have similar parameters as DMSO treated neurons (**B**) Mitochondrial length for inverse Aβ42 is comparable to DMSO treated neurons while Aβ42 oligomer treatment significantly reduces mitochondrial length. (**C**) Mitochondrial density for inverse Aβ42 is comparable to DMSO treated neurons while Aβ42 oligomer treatment significantly reduces mitochondrial density. **(D)** Spine density quantification for inverse Aβ42 is comparable to DMSO treated neurons while Aβ42 oligomer treatment significantly reduces spine density. All the analysis was done blind to the experimental conditions, and was done by manual counting using FIJI. Data is represented by box plots displaying minimum to maximum values, with the box denoting 25^th^, 50^th^ (median) and 75^th^ percentile from 3 independent experiments. n_INV_ = 41 dendrites, 243 mitochondria; n_veh_ = 31 dendrites; 178 mitochondria; n _Aβ42o_ = 33 dendrites, 245 mitochondria. Statistical analysis was performed using Kruskal-Wallis test followed by Dunn’s post-hoc test in (B-D). The test was considered significant when p<0.05 with the following criteria: * p<0.05; ** p<0.01; ***p<0.001; ****p<0.0001; ns, not significant. Scale bar= 2μm. (**E**) 16.5% Tris-Tricine SDS-PAGE was performed to resolve Aβ42 oligomers from the peptide monomer that were generated as described in Material and Methods. Immuno-blotting was performed with 6E10 antibody which shows a mixture of monomers, dimers and trimers in lane 1 and 2 (two independent oligomerization experiments). For quantifying the relative oligomer/monomer concentration in lane 1 and 2, Near-infrared fluorescence signal intensity was measured for dimer+trimer and monomer using an Odyssey Imager. Signal intensity values for Lane1: Dimer+Trimer-1510; monomer-1830. Signal intensity values for Lane2: Dimer+Trimer-736; monomer-1720.

**Figure S3.**
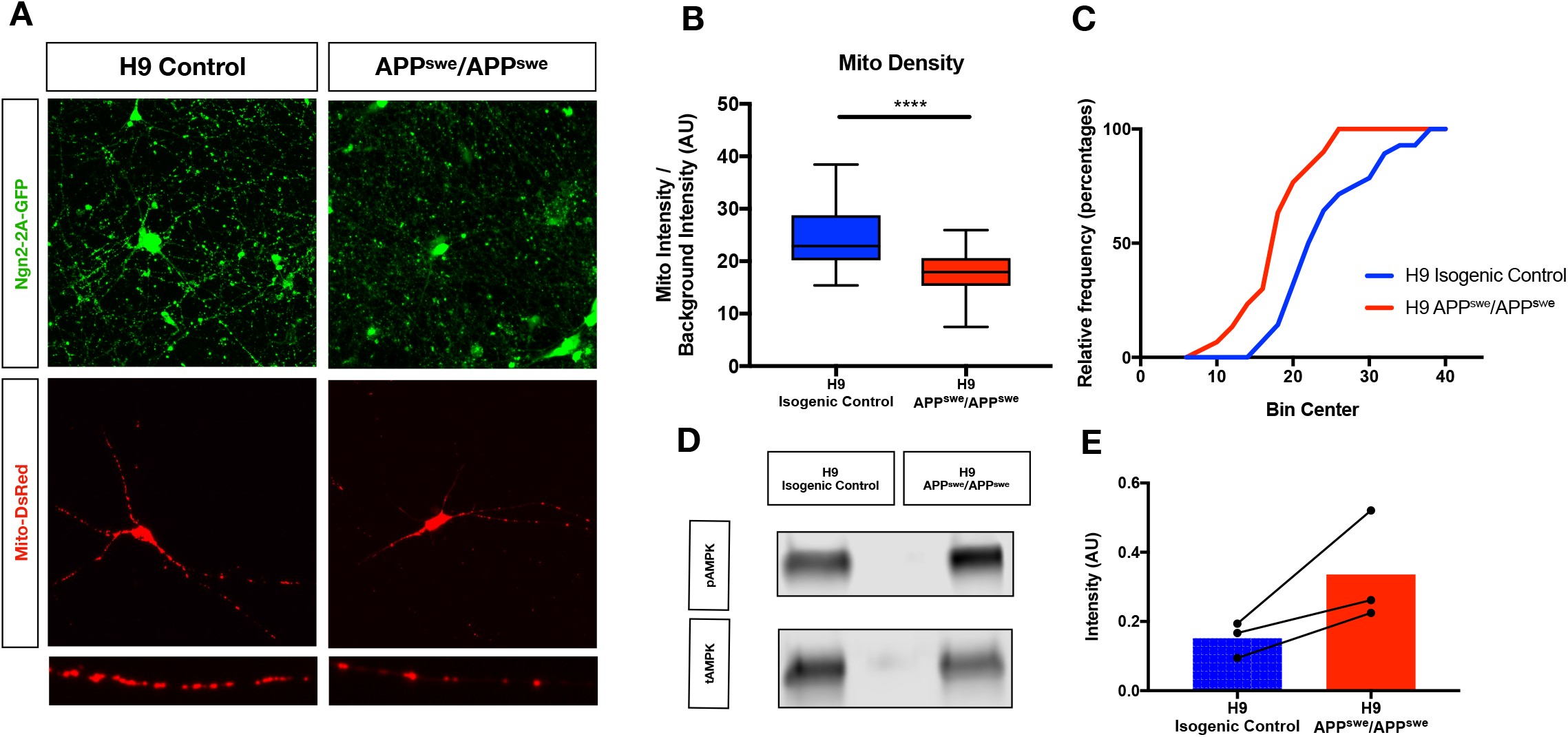
hESC-derived cortical neurons expressing endogenous APP^SWE^ mutations exhibit decreases in mitochondrial density and increased AMPK activation relative to isogenic controls. (**A**) Representative images neurons of control H9 embryonic stem (hESC) cells and H9: APP^SWE/SWE^ knockin hESCs at 24 DIV (post dox-induction). hESC-derived cortical-like neurons were induced using Ngn2-2A-eGFP driven neural induction (see methods), which allows visualization by eGFP. Neurons were transfected using Lipofectamine 3000 at 21-31 DIV with a pCAG-mitoDSRed reporter gene to label mitochondria, fixed 4 days post transfection, and imaged by confocal microscopy. Endogenous expression of the APP^SWE/SWE^ mutation is sufficient to induce a significant decrease in mitochondrial density in their dendrites (bottom high magnification panels) relative to isogenic control neurons. (**B**) Quantification of neuritic mitochondrial density. All of the analyses were done by kymographic fluorescent density measurement in Nikon Elements software. (**C**) Cumulative frequency distribution of mitochondrial density in Ngn2-induced cortical neurons derived from control H9 (blue) and APP^SWE/SWE^ cells (red). (**D**) Sample western blot of H9 isogenic control vs. H9: APP^SWE/SWE^ cortical neurons. 50μg of immunoprecipitated cell lysate was immunoblotted with phospho-AMPK (T172) and total AMPK to confirm activation of AMPK under endogenous expression of APP^SWE^ mutation. (**E**) Quantification of increased phospho-AMPK, representing mean and SEM. All of the statistical analyses were performed in Prism 6 (GraphPad Software). Data is represented by box plots displaying minimum to maximum values, with the box denoting 25th, 50th (median) and 75th percentile from 3 independent experiments. In A-C: n_H9 control_ = 227 neurites, 70 neurons; n_APPswe control_ = 203 neurites, 73 neurons. Statistical analyses were performed using One-way ANOVA followed by Kruskal-Wallis post-hoc test in B. The test was considered significant when p < 0.05 with the following criteria: **** p < 0.0001; ns, not significant.

**Figure S4.**
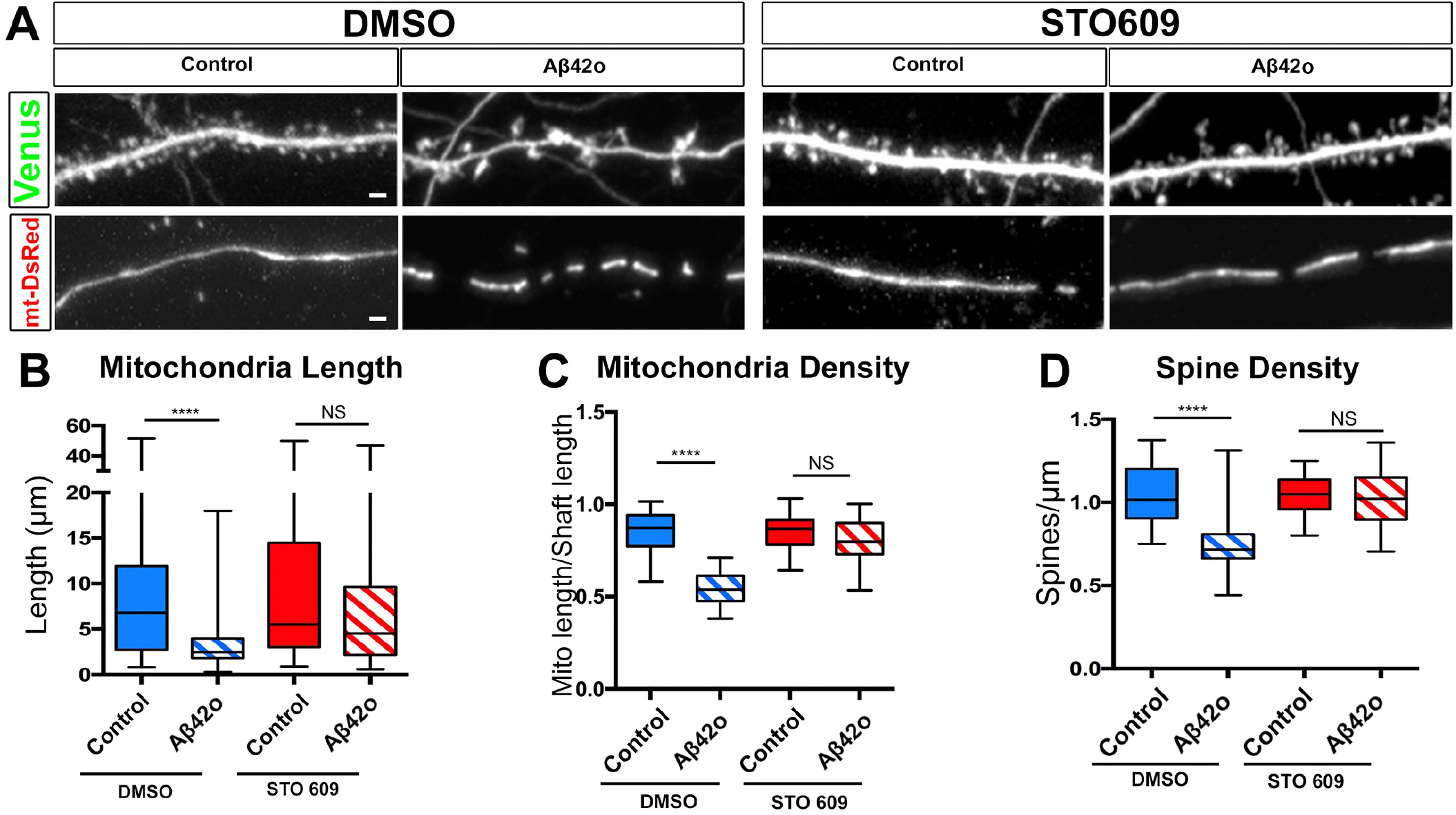
Oligomeric Aβ42 induced synaptotoxicity and dendritic mitochondrial fragmentation is CAMKK2 dependent. ***Related to Figure 4*** (A) Dendritic segments (secondary) of primary cortical neuron at 21 DIV. Embryos at E15.5 were *ex utero* electroporated with pCAG-Venus and pCAG-mito-DsRed. At 20 DIV, neurons were pre-treated with either DMSO or STO609 (2.5μM) for 2 hours before being treated with either control or Aβ42 oligomers for 24 hr. Blocking CAMKK2 activity by STO609 blocks both dendritic mitochondrial remodeling and synaptotoxicity. (B) Quantification of dendritic mitochondrial length, (C) dendritic mitochondrial density, and (D) spine Density. All the analysis was done blind to the experimental conditions, and was done by manual counting using FIJI. Data is represented by box plots displaying minimum to maximum values, with the box denoting 25^th^, 50^th^ (median) and 75^th^ percentile from 3 independent experiments. n_DMSO Control_ = 37 dendrites, 163 mitochondria; n_DMSO Aβ42o_ = 31 dendrites; 248 mitochondria; n_STO609 control_ = 37 dendrites; 163 mitochondria; n_STO609 Aβ42o_ = 35 dendrites, 157 mitochondria. All of the statistical analyses were performed with Prism 6 (GraphPad Software). Statistical analysis was performed using Kruskal-Wallis test followed by Dunn’s post-hoc test in (B-D). The test was considered significant when p<0.05 with the following criteria: * p<0.05; ** p<0.01; ***p<0.001; ****p<0.0001; ns, not significant. Scale bar= 2μm.

**Figure S5.**
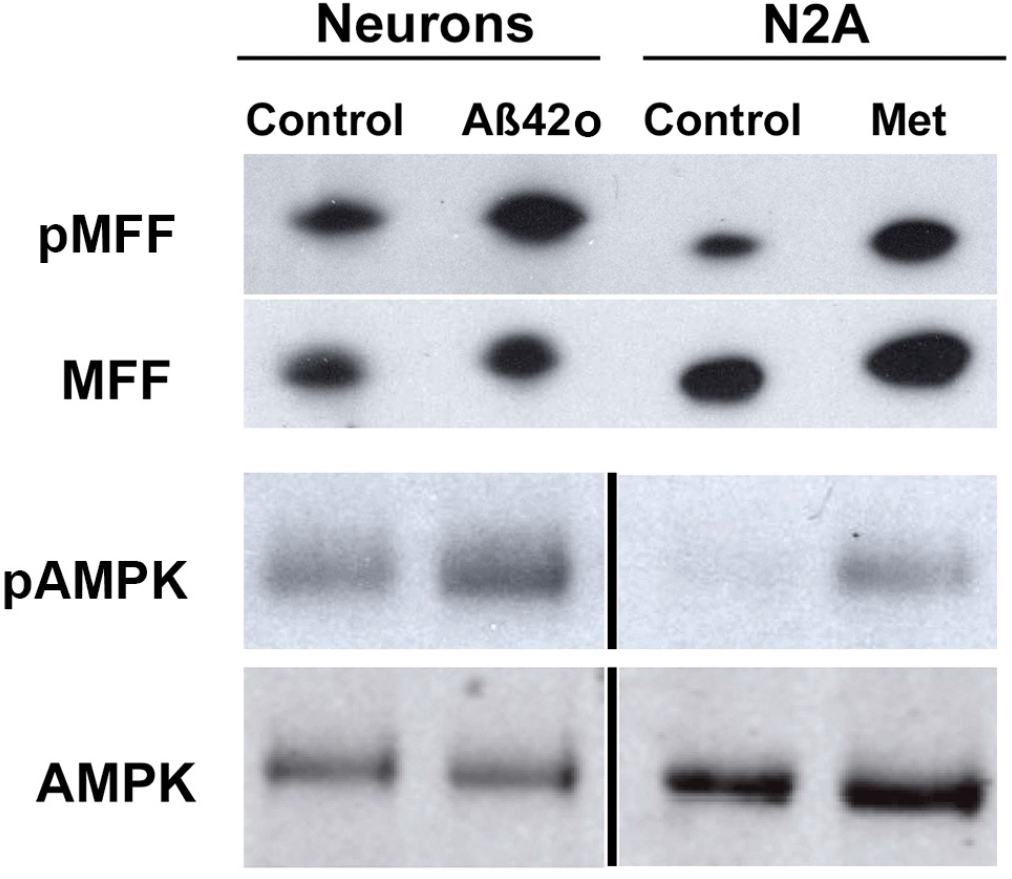
Aβ42o-dependent increase in MFF phosphorylation. ***Related to Figure 6*** Cortical PNs maintained in dissociated culture were treated at 20DIV with either control vehicle or Aβ42o for 24 hours. N2A cells were treated with either DMSO or 2mM of Metformin for 5 hours as positive control for AMPK-dependent MFF phosphorylation (Toyama et al., 2016). 80μg of whole cell lysate was immunoprecipitated with MFF antibody (Progen) and immunoblotted with MFF antibody or phospho-MFF antibody (S146, CST). Whole cell lysates were also immunoblotted with phospho-AMPK (T172, CST) and AMPK to confirm activation of AMPK under both Aβ42o for 24 hours (Mairet-Coello et al., 2013) and metformin treatment (Toyama et al., 2016).

**Figure S6.**
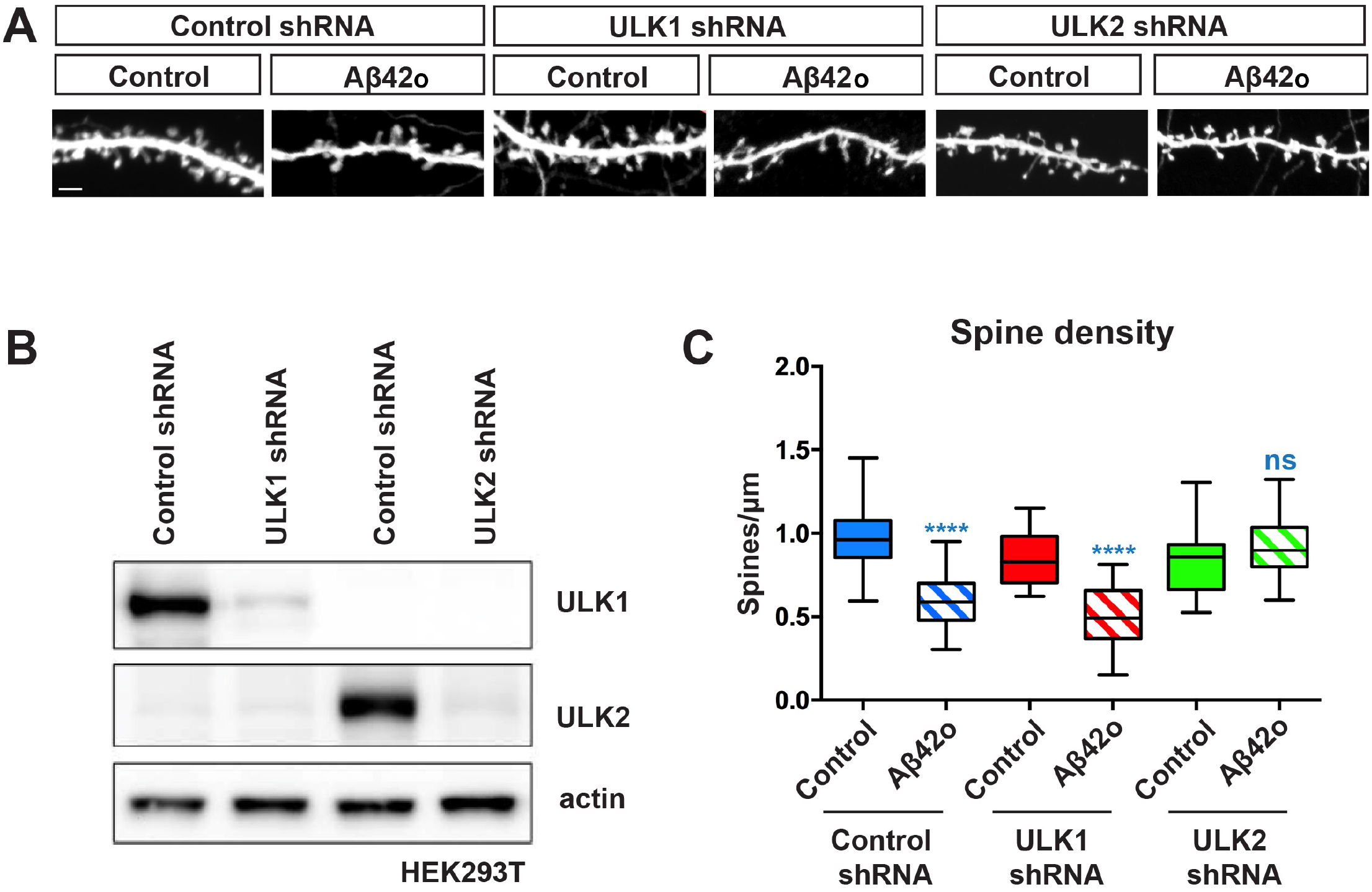
ULK2 but not ULK1 lies downstream is required for Aβ42o-dependent synaptotoxic effects. ***Related to Figure 7*** (**A**) Representative images of secondary dendritic segments of primary cortical neurons treated with either control or Aβ42o at 21DIV for 24 hours. Embryos were subjected to *ex-utero* electroporation at E15.5 with pCAG-Venus and control shRNA or an shRNA specific for the kinase ULK1 or ULK2. Knockdown of ULK2, but not ULK1 kinase blocks oligomeric Aβ42o-induced synaptotoxicity. (**B**) Western blot showing validation of ULK1 and ULK2 shRNA and the specificity of the shRNAs for the respective isoform. HEK 293T cells were transiently co-transfected with control shRNA or ULK1/ULK2 shRNA along with over-expressed myc-mULK1 and Flag-mULK2 respectively and western blotting was performed with 25μg of lysate with the indicated antibody. (**C**) Quantification of spine density for panel A. All the analysis was done blind to the experimental conditions, and was done by manual counting using FIJI. All of the statistical analyses were performed with Prism 6 (GraphPad Software). Data is represented by box plots displaying minimum to maximum values, with the box denoting 25^th^, 50^th^ (median) and 75^th^ percentile from 3 independent experiments. n_PLKO control_ = 31 dendrites; n_PLKO Aβ42o_ = 27 dendrites; n_ULK1shRNA control_ = 27 dendrites; n_ULK1shRNA Aβ42o_ = 24 dendrites; n_ULK2shRNA control_ = 33 dendrites; n_ULK2shRNA Aβ42o_ = 35 dendrites. Statistical analysis was performed using Kruskal-Wallis test followed by Dunn’s post-hoc test in (C). The test was considered significant when p<0.05 with the following criteria: * p<0.05; ** p<0.01; ns, not significant. Scale bar= 2μm.

**Figure S7.**
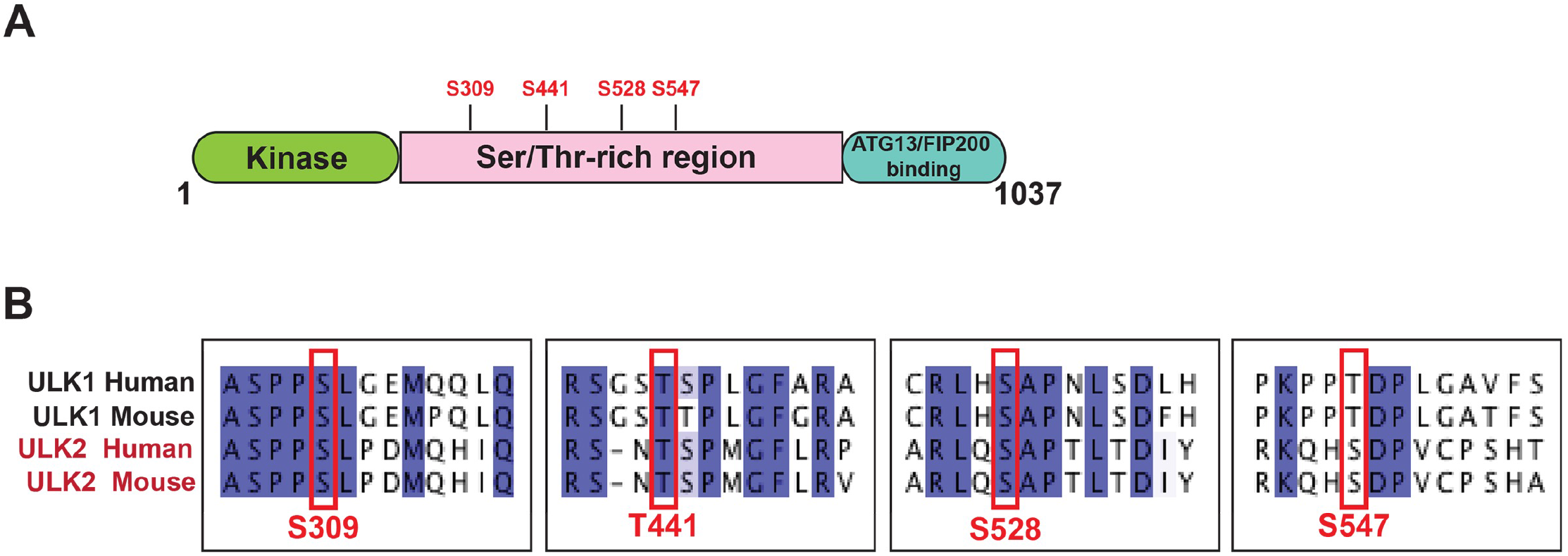
Schematic of conserved AMPK-mediated phosphorylation sites in ULK2. ***Related to Figure 8*** Schematic of mULK2 domain structure highlighting the four predicted AMPK-mediated phosphorylation sites. (B) ClustalW multiple sequence alignment of human and mouse ULK1 and ULK2 showing a high degree of conservation of four AMPK-mediated phosphorylation sites in ULK2 which were previously reported in ULK1 (Toyama et al., 2016).

**Supplementary Movie S1** Time-lapse of DIV 21 Inverse peptide treated cortical pyramidal neurons expressing mito-mTagBFP2, LAMP1-mEmerald and mRFP-LC3 to visualize mitochondria, lysosome and autophagosome (dendritic segment corresponds to the representative image in Fig. 3). Neurons were imaged every 15min for 14hr. *Related to Figure 3*.

**Supplementary Movie S2** Time-lapse of DIV 21 Aβ42o treated cortical pyramidal neurons expressing mito-mTagBFP2, LAMP1-mEmerald and RFP-LC3 to visualize mitochondria, lysosome and autophagosome (dendritic segment corresponds to the representative image in Fig. 3). Neurons were imaged every 15min for 14hr. *Related to Figure 3*.

## Supplementary Experimental Procedures

### Constructs

pSCV2-Venus and pCAG-mito-dsRed were described in previous publication (Lewis et al., 2016). pLKO-MFFsh specific for mouse was obtained from Sigma Aldrich (TRCN0000174665). Validation of the shRNA knockdown efficiency was done elsewhere (Lewis et al., 2018). cDNA encoding Renalin, MFF WT (Uniprot Q9GZY8 isoform 5), and MFF AA were described previously (Toyama et al., 2016) and sublconed in the pCAG plasmid backbone 3’ to CAG promoter using PCR. 0.25μg/mL of individual constructs were used for the MFF rescue experiments. pCAG-Cre, pCAG-Scrambled Cre, pCIG-hTau (isoform 4R2N), and non-phosphorylatable hTau-S262A were described in previous publications (Courchet et al., 2013). pcDNA3-flag-ULK2 specific for mouse was obtained from Addgene (#27637) and was used to generate the kinase-inactive (K39I) and serine-to-alanine 4SA mutant (S309/T441/S528/S547A) using Quikchange II site directed mutagenesis kit (Agilent technologies) and were subcloned into pCAG vector backbone using PCR. pSUPER-scrambled, ULK1 shRNA and ULK2 shRNA constructs were validated in a previous study (Zhou et al., 2007).

### *In utero* hippocampal electroporation

Timed pregnant hybrid F1 females were obtained by mating inbred 129/SvJ females and C57Bl/6J males. These F1 females were then mated with J20 males to obtain both mice that do express the APP transgene and mice that do not express the transgene in the same littermate. *In utero* electroporations were performed as detailed in (Szczurkowska et al., 2016) with the exception that J20 mice were used. Briefly, a mix of endotoxin-free plasmid preparation (PSCV2-mVenus −1μg/mL, pCAG-mitoDsRed −0.5 μg/mL) was injected into both lateral hemisphere of E15.5 embryos using a picospritzer. Electroporation was performed with triple electrode to target hippocampal progenitors in E15.5 embryos by placing the two anodes (positively charged) on either side of the head and the cathode on top of the head at a 0° angle to the horizontal plane. Four pulses of 45V for 50ms with 500ms interval were used for the electroporation.

### Tissue slice preparation and Immunofluorescence

Animals were sacrificed 3 months after birth by terminal transcardial perfusion of 4% paraformaldehyde (PFA, Electron Microscopy Sciences) followed by overnight post-fixation in 4% PFA. Brains were washed 3X with PBS for 5 minutes and sectioned at 100μm thickness using vibratome (Leica). Sections were permeabilized with 0.2% Triton X-100 3X for 5 minutes, blocked in PBS based blocking buffer with 5% BSA and 0.2% Triton, and stained with chicken anti-GFP (Aves, 1:1000) and rabbit anti-RFP (Abcam, 1:1000) for enhancement of PSCV2-mVenus and pCAG-mitoDsRed, respectively overnight at 4°C. Sections were washed with 0.2% Triton buffer 3X for 5 minutes and incubated with Alexa488- and Alexa 555-labeled secondary antibodies for PSCV2-mVenus and pCAG-mitoDsRed, respectively overnight at 4°C. Sections were washed with 0.2% Triton buffer 3X for 10 minutes each and mounted using VectaShield® Mounting Medium (Vector Laboratory).

### Drug Treatments

Neuronal cells at 20-25 DIV were treated with STO609 (2.5μM, Millipore), a CAMKK2 inhibitor, 2.5 hours prior to Aβ42 treatment. N2A cells were transfected with pCAG-MFF-WT construct using the recommended protocol from FuGENE®HD Transfection Reagent overnight. Cells were treated with either DMSO or Metformin (2mM, Sigma) for 5 hours before cells were lysed using Triton lysis buffer as described under Cell Lysis and Immunoprecipitation.

### eSC Immunoprecipitation and Western Blotting

For phospho-T172 AMPK and total AMPK for eSC experiments, equal amounts of lysates (50µg) were immunoprecipitated with phospho-T172 AMPKα or total AMPKα to enhance AMPK proteins. Lysates from H9-Control and H9APP lines were incubated overnight at 4°C in a rotator with either phospho-T172 AMPK or total AMPK primary antibodies. Protein A-Agarose beads (Sigma) were washed three times with PBS and added to the immuno-bound lysates for at least 1 hr at 4°C. The lysates and beads were then washed thrice with lysis buffer and eluted in SDS lysis buffer for 5 minutes at 95°C and resolved by loading equal amounts of eluted protein on a Mini-Protean TGX (4-20%) SDS-PAGE (Bio-rad). The separated proteins were transferred onto polyvinylidene difluoride membrane (PVDF, Bio-Rad), and blocked for 1 hour with Odyssey Blocking Buffer (PBS). Membranes were then incubated overnight at 4°C with phospho-T172 AMPK or total AMPK primary antibodies diluted in the same blocking buffer. Incubations with Li-Cor fluorescence-coupled secondary antibodies were performed for 1 hour at room temperature, and visualization was performed by Li-Cor Odyssey Blot Imager.

### Analysis of Spine Density and Mitochondrial Morphology

Dendritic spine densities were estimated on secondary or tertiary dendritic branches for cultured neurons and in the depth of the z stack for slices, using FiJi software (ImageJ, NIH). The length of the dendritic segment was measured on the z projection, which implies the density could be overestimated. To limit this issue, only dendrites that were parallel to the plane of the slices were analyzed. Spines were quantified over an average of 40μm in cultures (between 2-3 segments per cell) and 40μm *in vivo* (between 2-3 segments per cell). Spine density was defined as the number of quantified spines divided by the length over which the spines were quantified. The criteria for measuring mitochondrial length for each dendritic segment were same as above. Mitochondrial density was defined as the total sum of all the mitochondria divided by the length over which the mitochondria lengths were measured.

### Analysis of Mitophagy Events

For Fig.2B, manual counting of total number of dendrites and number of dendrites showing accumulation of fluorescently-tagged LC3-RFP and LAMP1-mEmerald was performed for all imaged neurons. To confirm both LC3 and LAMP1 accumulation over time following Aβ42o application using an independent quantification approach, we selectively examined all of the 251 dendritic segments categorized as showing LC3+/LAMP1+ accumulation upon Aβ42o application and measured the intensities of LC3 and LAMP1 fluorescence (within an ROI ranging from 20-30μm in length) over time. All the dendrites that showed accumulation of LC3 and LAMP1 in control conditions were also analyzed, as well as randomly selected dendritic segments that did not show detectable accumulation of LC3 and LAMP1. Optical density of LAMP1-mEmerald and LC3-RFP fluorescence were normalized to the initial starting fluorescence intensities at t_0_ (ΔF/F_0_). For these selected dendrites, mito-mtagBFP2 intensities were also measured over time for Fig. 2E. To measure local mitophagy events as indicated in Fig. 2F, random dendrites analyzed in Fig.2B were selected for each condition. For a given dendritic segment, intensity line scan analysis (pixel neighborhood-1 pixel) was performed where two vertical lines were drawn: one along the region of dendrite without LC3/LAMP1 puncta and one along the region with LC3/LAMP1 puncta. The fold change of mitochondrial intensity from time point zero to 14hr after respective treatment was calculated as illustrated in Fig. 2F.

### Analysis of Mitochondrial Density in hES cell-derived cortical neurons

For Figure S7B-C, lines of equal thickness and of an average length of ∼40μm were drawn along the neurites of differentiated hES cell-derived cortical neurons in Nikon NIS-Elements Software. A kymograph was subsequently created from each drawn line, for which the mean fluorescence intensity of the mitochondria as a function of area was calculated and normalized to the background fluorescence intensity of the image.

